# Recording Temporal Signals with Minutes Resolution Using Enzymatic DNA Synthesis

**DOI:** 10.1101/2021.07.14.452380

**Authors:** Namita Bhan, Alec Callisto, Jonathan Strutz, Joshua Glaser, Reza Kalhor, Edward Boyden, George Church, Konrad Kording, Keith E.J. Tyo

## Abstract

Employing DNA as a high-density data storage medium has paved the way for next-generation digital storage and biosensing technologies. However, the multipart architecture of current DNA-based recording techniques renders them inherently slow and incapable of recording fluctuating signals with sub-hour frequencies. To address this limitation, we developed a simplified system employing a single enzyme, terminal deoxynucleotidyl transferase (TdT), to transduce environmental signals into DNA. TdT adds nucleotides to the 3’ ends of single-stranded DNA (ssDNA) in a template-independent manner, selecting bases according to inherent preferences and environmental conditions. By characterizing TdT nucleotide selectivity under different conditions, we show that TdT can encode various physiologically relevant signals like Co^2+^, Ca^2+^, Zn^2+^ concentrations and temperature changes *in vitro*. Further, by considering the average rate of nucleotide incorporation, we show that the resulting ssDNA functions as a molecular ticker tape. With this method we accurately encode a temporal record of fluctuations in Co^2+^ concentration to within 1 minute over a 60-minute period. Finally, we engineer TdT to allosterically turn off in the presence of physiologically relevant concentration of calcium. We use this engineered TdT in concert with a reference TdT to develop a two-polymerase system capable of recording a single step change in Ca^2+^ signal to within 1 minute over a 60-minute period. This work expands the repertoire of DNA-based recording techniques by developing a novel DNA synthesis-based system that can record temporal environmental signals into DNA with minutes resolution.

## Main Text

### Introduction

DNA is an attractive medium for both long-term data storage and for *in vitro* recording molecular events due to its high information density (1–3) and long-term stability (4). Molecular recording strategies write information into DNA by altering existing DNA sequences (5) or adding new sequences (6). For example, systems have been developed that use methods including differential CRISPR spacer acquisition (5, 7, 8), enzymatic synthesis (9–11) and others (1, 8, 12). By connecting these DNA modifications to a user input (in the case of data storage) or environmental signal of interest (in the case of recording events), these strategies enable *post hoc* recovery of signal dynamics over time by DNA sequencing. To date, molecular recording systems, both *in vitro* and *in vivo*, have connected signals of interest to DNA recordings with transcriptional control, using signal-responsive promoters to drive the expression of molecular writers, such as base-editors, CRISPR-associated systems, or gene-circuits, to record changes in signal. These approaches have yielded accurate recordings, however the time required to transduce signals through a recording apparatus that includes transcription, translation, and DNA modification fundamentally constrains the application of these methods to events on the timescales of hours or days. A recording mechanism that relies only on post-translational elements would be inherently faster as signal transduction would only require sub-second conformational shifts in one enzyme.

In an effort to speed up DNA recording processes, we hypothesized that a DNA polymerase (DNAp), which continually incorporates bases (13), could serve as a candidate for post-translational molecular encoding. In such a system, a DNAp functions as a “ticker-tape” recorder, transforming changes in environmental signals into changes in the composition of the DNA it synthesizes (14) (Fig. 1A). Much faster than transcription and translation, nucleotide incorporation occurs on a timescale on the order of milliseconds to seconds (15), potentially enabling orders of magnitude improvements in the temporal accuracy and resolution of molecular recording. However, prototypical DNAp’s replicate the contents of an existing strand, which would prevent recording of new information. A DNAp that does not simply replicate DNA but rather creates a *de novo* sequence could allow for DNA recording.

**Figure 1:**
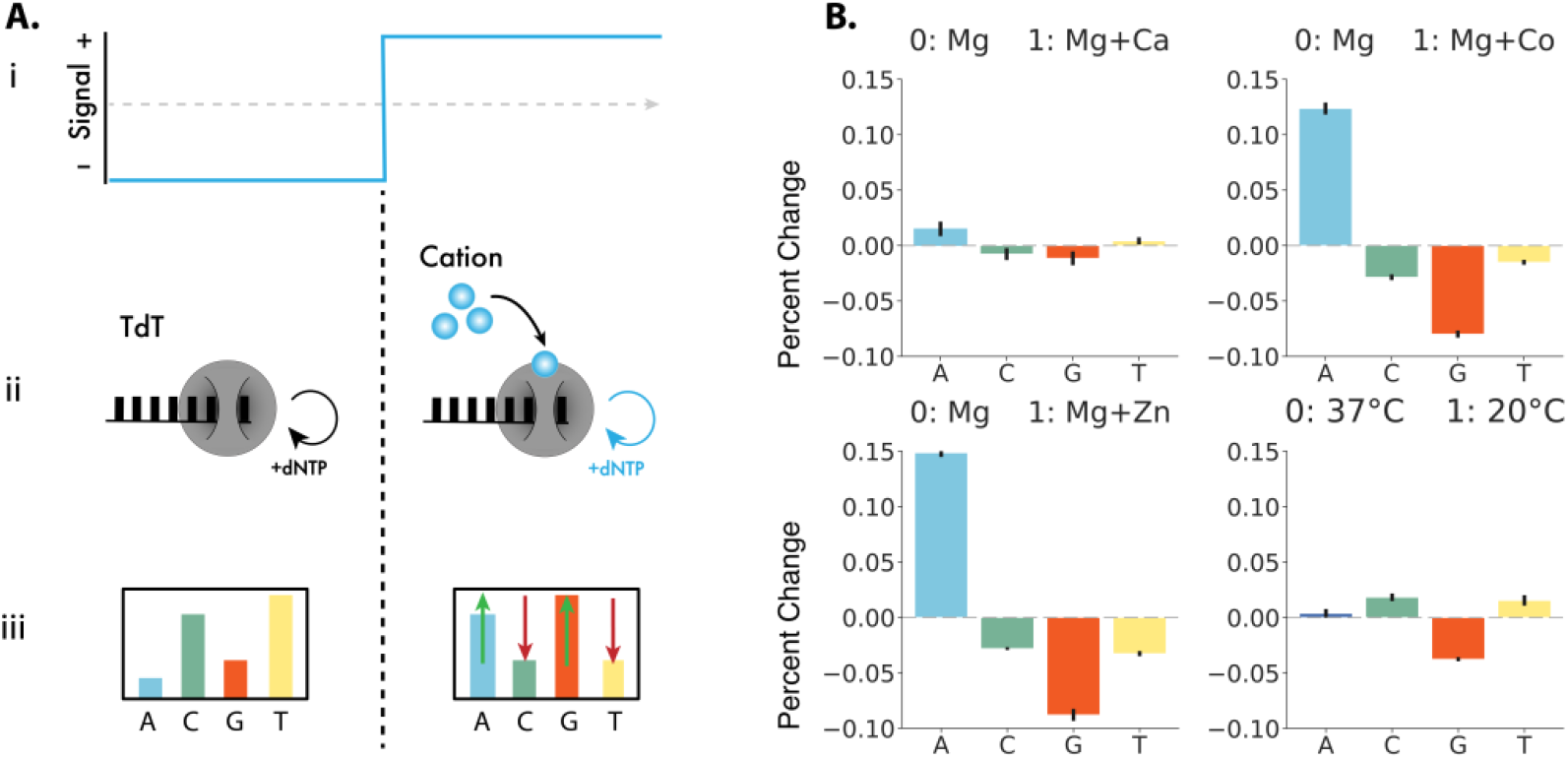
TURTLES device architecture and environmental signal responses. **(A.i)** A representative time-varying input signal. **(A.ii)** TdT interacts directly with the signal of interest (small blue circles) **(A.iii)** resulting in different average DNA compositions in each condition. **(B)** Change in frequency of nucleotide selectivity by TdT in the presence of various environmental signals tested. Signal 0 is 10 mM Mg^2+^ at 37 °C for 1 hour. Signal 1 was: (top-right) 10 mM Mg^2+^ + 1 mM Ca^2+^ at 37 °C for 1 hour A increased by 1.5%, G decrease by 1.2%, T increased by 0.4%, and C decreased by 0.8%; (top-left) 10 mM Mg^2+^ + 0.25 mM Co^2+^ at 37 °C for 1 hour, A incorporation increased by 12.4%, while G decreased by 8.0% and T and C decreased by 1.5% and 2.9% respectively; (bottom-left) 10 mM Mg + 20 μM Zn^2+^ at 37 °C for 1 hour, A increased by 14.9%, G 8.8% decreased by, T decreased by 3.3%, and C decreased by 2.8%; and (bottom-right) 10 mM Mg^2+^ at 20 °C for 1 hour, 0.4% increase in A, 3.8% decrease in G, 1.5 % increase in T and 1.8% increase in incorporation of C. Error bars show two standard deviations of the mean. Statistical significance was assessed after first transforming the data into Aitchison space which makes each dNTP frequency change statistically independent of the others (Fig. S2).

Terminal deoxynucleotidyl transferase (TdT) is a DNAp that can randomly incorporate bases to the 3’ of a DNA strand with biases toward particular bases (16, 17). Shifting the nucleotide bias of TdT could make it a prime candidate for post-translational control of DNA encoding. In fact, *in vitro* experiments have shown cations (e.g., Co^2+^) can shift the bias of TdT (16, 18). In addition, DNA is synthesized in a sequential manner which provides an estimate of the time a particular base is added. We therefore hypothesized that the environment in which a TdT extends a DNA strand might be encoded by the average base-composition of the extended DNA. Put another way, by combining the change in nucleotide bias in the presence of cations and the time bases are added inferred from sequence, a molecular ticker tape may be possible (13, 14).

Here, we introduce TdT-based Untemplated Recording of Temporal Local Environmental Signals (TURTLES), a polymerase-based molecular recording system that achieves high time resolution *in vitro* by utilizing post-translational control to change the bases incorporated. First, we describe methods to characterize DNA sequences synthesized by TdT and show that cation concentrations can be encoded in populations of TdT-synthesized DNA using an approach that analyzes the average composition of several bases added at similar times on the same or parallel strands of DNA. We next developed an algorithm to accurately estimate the times of signal changes and show that temporal information can be accurately recovered by using estimates of DNA synthesis rates to map DNA sequences to real time. We also describe an expanded TURTLES system that uses an engineered, allosterically modulated TdT to expand the generalizability and tunability of the system. By inserting an exogenous sensing domain, we show that TURTLES can be adapted to arbitrary signals of interest. Taken together, these results establish the feasibility of DNA synthesis-based encoding systems and demonstrate minutes resolution recording of cationic environmental signals for enhanced applications in DNA data storage and DNA recording.

## Results

### TdT can encode environmental signals *in vitro* via changes in base selectivity

The cations present in the reaction environment of TdT affect the rate of incorporation for specific nucleotides (16). For example, previous studies (18–20) and our experiments show that when only one nucleotide is present, the incorporation rates of pyrimidines, dCTP and dTTP, increase in the presence of Co^2+^ (Fig. S1).

We sought to examine if these Co^2+^-dependent changes in kinetics also occurred in the presence of all four nucleotides, dATP, dCTP, dGTP, and dTTP (hereafter referred to as A, C, G, and T). The nucleotide composition of ssDNA extended by bovine TdT in a Cobalt-free reaction buffer or with cobalt added was determined by next generation sequencing. In the presences of Co^2+^, A incorporation increased, while G, T and C incorporation decreased (Fig. 1B and Fig. S2). Notably, the significant difference in the composition of DNA each condition effectively encodes information about the environmental Co^2+^ concentration at the time of DNA synthesis.

Next, we were interested in understanding which conditions could be encoded by TURTLES. We examined Ca^2+^, Zn^2+^, and temperature. Ca^2+^ signaling is biologically ubiquitous and functions in neural firing (21), fertilization (22, 23), and neurodevelopment (24); Zn^2+^ is an important signal in the development and differentiation of cells (25); and temperature is relevant in many situations.

Each signal altered both the particular dNTPs affected and the magnitude of the change in dNTP selectivity. We were able to encode 20 μM Zn^2+^, 1 mM Ca^2+^ and the temperature of 20 °C (Fig. 1B and Fig. S2). Both cation addition and temperature change also altered the lengths of ssDNA strands synthesized (Fig. S3-8). For each environmental condition tested, we observed significant differences in the composition of TdT synthesized DNA. We conclude that input-dependent changes in TdT nucleotide selectivity can encode environmental information into DNA. For further analysis we chose to focus on Co^2+^ as the candidate cationic signal due to the large difference in TdT selectivity.

### Recording a single step change in Co^2+^ concentration with minutes resolution

Having shown nucleotide selectivity changes in the presence of Co^2+^, we attempted to identify the time at which Co^2+^ was added to a TdT-catalyzed ssDNA synthesis reaction based on the changes in the nucleotide composition of the synthesized ssDNA (Fig. 2A). During a 60-minute extension reaction, we created step transitions in cobalt concentration by adding 0.25 mM Co^2+^ at 10, 20, and 45 minutes (hereafter referred to as a 0→1 input where ‘0’ is Co^2+^-free and ‘1’ is with 0.25 mM added Co^2+^) (Fig. 2B top). For each reaction, we analyzed approximately 500,000 DNA strands by deep sequencing and calculated the dNTP incorporation frequencies over all reads. Because the change in dNTP selectivity is a compositional data type (i.e., all changes in base frequency sum to 0%), they are not independent and do not satisfy the independence assumption required for most statistical tests. Therefore, to perform hypothesis testing, base composition was transformed into Aitchison space, where the proportion of each base becomes independent of the other three. The output signal for each reaction was calculated as the normalized distance in Aitchison space between the 0 and 1 controls.

**Figure 2:**
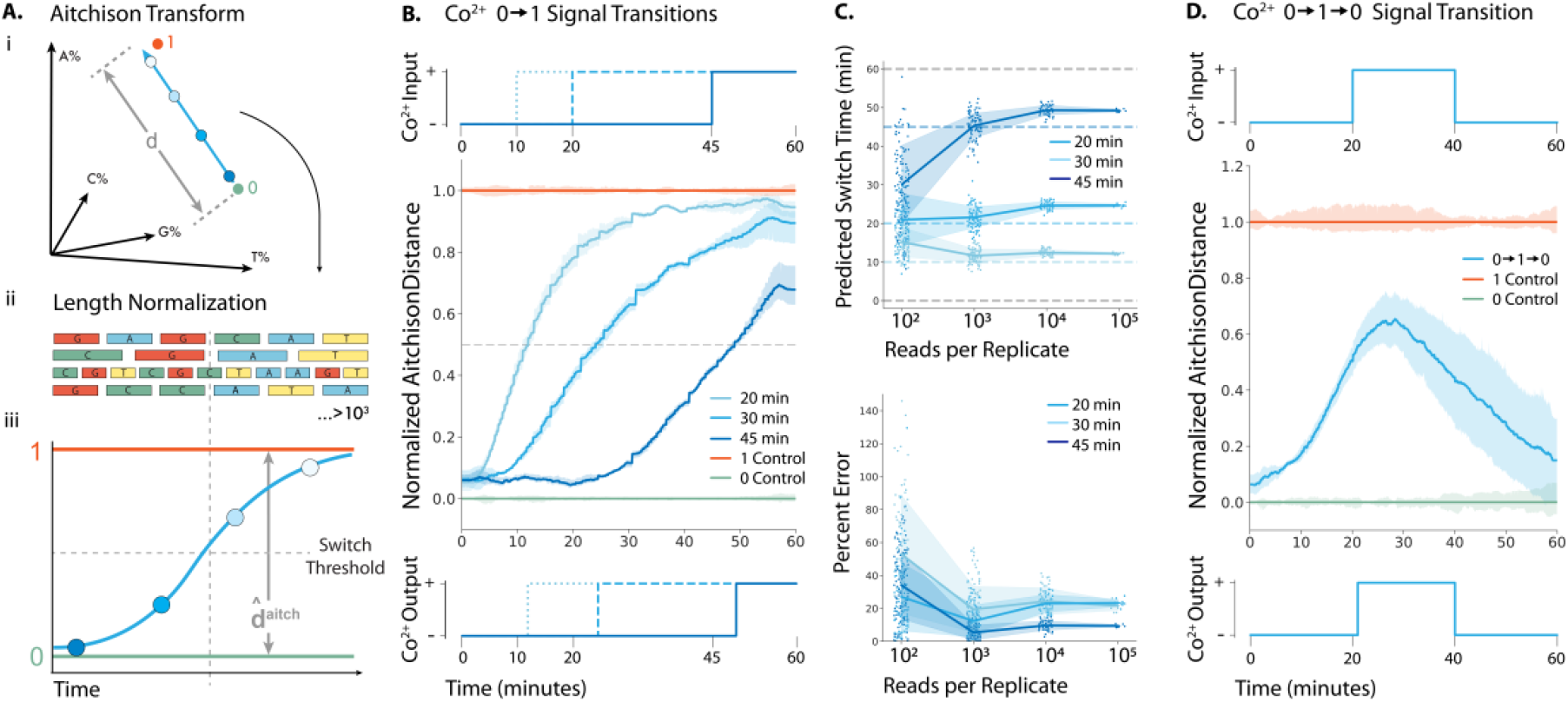
Recording a Co^2+^ fluctuations into ssDNA with minutes resolution *in vitro*. **(A.i.)** Representation of how percent incorporation of each nucleotide is dependent on each of the nucleotide incorporated. **(A.ii.)** Sequences were normalized by length before the nucleotide composition at each time point was calculated. **(A.iii.)** By transforming the percent incorporation of each nucleotide to the Atichison distance we can calculate the total “output signal”. We plot the Aitchison distance for the recording experiment between the 0 (green) and 1 (orange) signal. **(B) Top:** 0.25 mM Co^2+^ was added at time 10, 20 and 40 minutes to generate a 0→1 transition. **Center:** Mean output signal across 3 biological replicates. Vertical lines are drawn at the inferred transition time. **Bottom:** Predicted output signal transition times were 11.9, 24.4 and 49.2 minutes. **(C) Top:** Predicted switch times for each 0→1 transition calculated from randomly sampled subsets of sequences. **Bottom:** Time prediction error for each 0→1 transition calculated from randomly sampled subsets of sequences. **(D) Top:** 0.25 mM Co^2+^ was added at 20 minutes and then removed at 40 minutes to generate a 0→1→0 transition. **Center:** Mean output signal across 3 biological replicates. **Bottom:** Using the algorithm detailed in Glaser et al. (13), the signal was deconvoluted into a binary response, with vertical lines drawn at the predicted switch times of 21 minutes and 41 minutes.

After normalizing each sequence by its own length, the Aitchison location along the extended strands showed that later addition of Co^2+^ resulted in dNTP selectivity changes farther down the extended strand (Fig. 2B center). To estimate the real time at which changes occurred, we calculated the average location along the strand for all sequences in each condition at which a distance halfway between the 0 and 1 control output signal was reached. To translate this location into a particular time in the experiment, we calculated the average rate of dNTP addition in each state (Fig. S9) and derived an equation that adjusted for the change in rate of DNA synthesis between the 0 and 1 controls (Equation 5, Supplementary Methods 3). Using this information, we estimated that the Co^2+^ additions were made at 11.9, 24.4 and 49.2 minutes (Fig. 2B bottom). We were also able to estimate the time within 4 minutes of the unit input step function for the reverse, a 1→ 0 condition (Fig. S10 &11).

While we were able to accurately estimate the times of Co^2+^ addition (0→1) and removal (1→0), simultaneously synthesizing ~500,000 strands of DNA will be infeasible for certain applications. To determine the number of strands needed for reasonable statistical certainty, we randomly sampled smaller groups of strands from the experiment and evaluated our ability to predict when Co^2+^ was added (Fig. 2C top and bottom). With about 1,000 strands, we still estimated the time of Co^2+^ additions to within 2 minutes of actual input times (Fig 2C,). Thus, TURTLES-1 recordings are robust and encode high resolution temporal information even with a limited number of ssDNA substrates.

### Recording multiple fluctuations in Co^2+^ concentration onto DNA with minutes resolution

In contrast to current DNA-based recorders, which rely on time-integrated recording methods (i.e., accumulation of mutations) or slow signal transducing steps, DNA synthesis-based approaches can record the dynamics of multiple fluctuations in real time. While accumulation can tell what fraction of the time a signal was present in a time period, the ability to record multiple temporal changes would enable new levels of insight into dynamical processes such as physiologically signaling, which are poorly captured by time-integrated recording methods.

We used TURTLES to record a 0→1→0 input cobalt signal. The 0 condition was maintained first 20 minutes, 1 for the next 20 minutes, and 0 for the last 20 minutes of the extension reaction (Fig. 2D top). Using the same methods as the single step transition, we calculated the output signal (Fig. 2D center). To account for the additional complexity of multiple fluctuations, we used an algorithm previously developed by Glaser *et al*.(13) (see Materials and Methods for details) to binarize the value of the output signal every 0.1 min. We were able to accurately reconstruct the input 0→1→0 signal, estimating transitions between the 0 and 1 signals occurring at 21 and 41 minutes based on sequencing data (Fig. 2D bottom).

Based on the measured experimental parameters, we used *in silico* simulations to estimate the performance of TURTLES in more complex recording environments. We investigated how rapidly signals could change and still be detected and how many consecutive condition changes could be recorded accurately. By varying the length of time of each input condition (0 or 1) from 1 to 20 minutes (Fig. S12A), we estimated that TURTLES can record 6 consecutive signal changes with 1 minute between each with >75% accuracy from ≥ 2000 strands of 100bp ssDNA synthesized (>90% accuracy with ≥ 60,000 strands of ssDNA of 50 bp length each) (Fig. S12B). By keeping the duration of each input condition (0 or 1) constant at 10 minutes (Fig S12C) and varying the total number of condition changes, we estimated that TURTLES would be capable of recording 10 sequential input signal changes with >80% accuracy (Fig. S12D). We thus show that TURTLES has unprecedented temporal precision and can robustly decode signals across a range of frequencies.

Having successfully encoded three sequential environmental states with TURTLES-1, we considered the feasibility of TURTLES-based digital data encoding. In this context, the data density of TURTLES-1 recordings is determined by the number of inputs that yield distinguishable outputs. By altering the ratios of nucleotides provided as substrates to a reaction in sequential steps, the composition of DNA synthesized by TdT could be controlled with high resolution. Assuming 5% changes in nucleotide composition can be distinguished, up to 11 bits of data can be encoded in each step for 33 bits in a three-step recording. The bits encoded in each step would increase to 17 bits if 1% changes can be distinguished (Fig S13).

### TdT can be engineered to allosterically respond to and encode environmental signals

Unlike Co^2+^ and Zn^2+^, we observed that Ca^2+^ only modestly altered the dNTP selectivity of TdT, precluding temporal recordings of Ca^2+^ concentration. To show that TURTLES could be expanded to signals to which TdT was unresponsive or weakly responsive, we attempted to engineer a TdT to allosterically respond to Ca^2+^. The structural determinants of base selectivity in TdT are poorly understood, which ruled out directly increasing the dynamic range of Ca^2+^-responsive dNTP selectivity changes (26). Accordingly, we conceived a modular recording system based on two distinct TdT species with different inherent dNTP selectivity.

The two-TdT system, TURTLES-2, uses a reference TdT whose catalytic rate is unaffected by inputs and a sensor TdT that is allosterically activated or deactivated in response to input signals. By choosing a pair of sensor and reference TdTs with distinct nucleotide selectivity, TURTLES-2 encodes environmental signals into changes in DNA composition based on the differential activity of the two TdTs (figure 3A). As the sensing and recording functions of the system are distributed between two TdT variants, TURTLES-2 is more accessible to tuning and engineering efforts.

**Figure 3:**
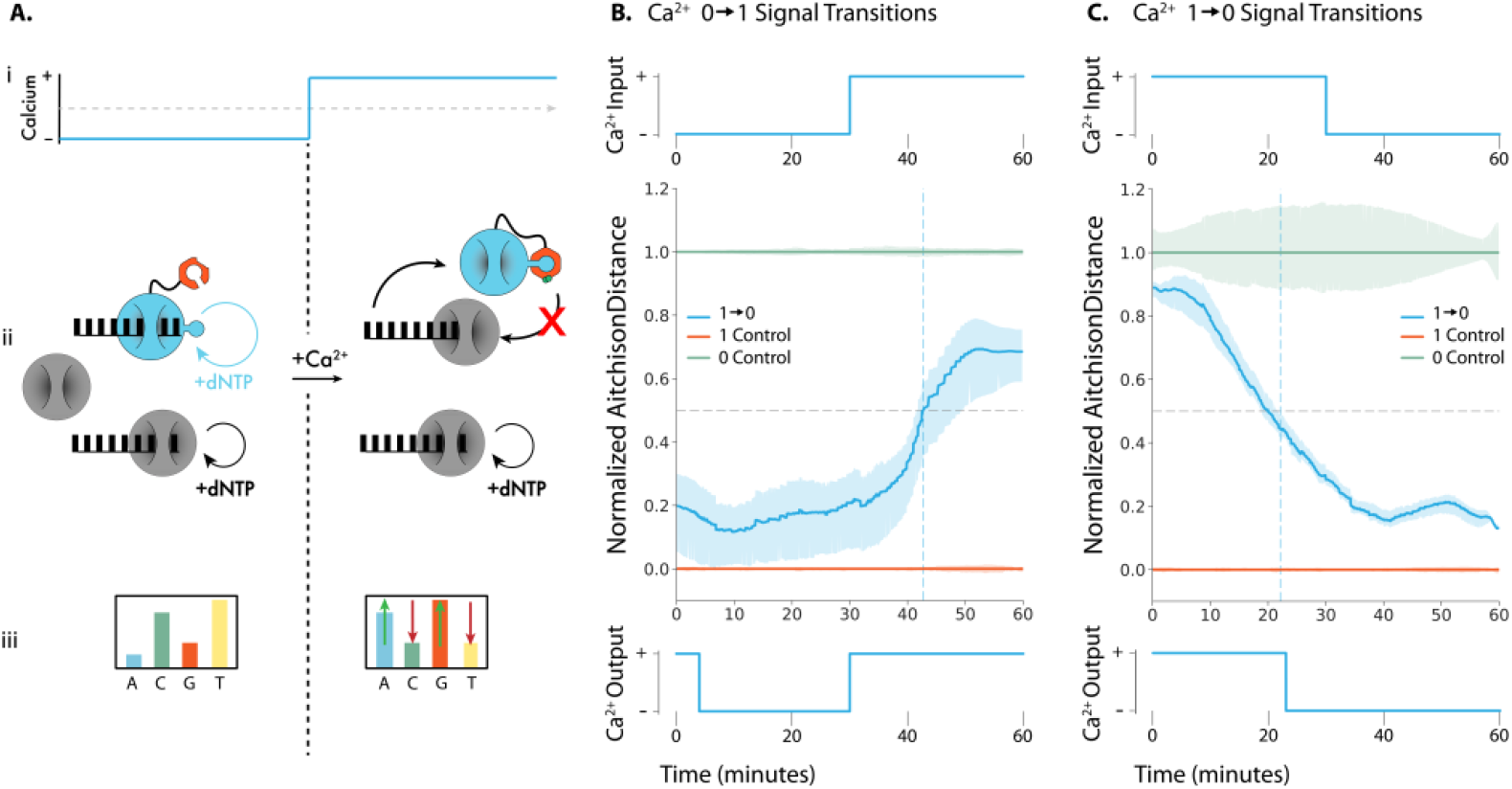
Recording Ca^2+^ changes into ssDNA with TURTLES-2. **(A.i.)** Representative calcium transition changing concentration from calcium-free to calcium-added conditions during a TdT-based DNA synthesis reaction. Mg^2+^ concentration and reaction temperature are held constant. **(A.ii.)** The CaM subunit (orange) of engineered TdT (teal) binds with the fused M13 peptide in the presence of Ca^2+^ to allosterically turn off DNA synthesis. The activity of the reference TdT (grey) is not affected by Ca^2+^. In the absence of Ca^2+^ both the engineered TdT and reference TdT are carrying out DNA synthesis. In the presence of Ca^2+^ only the reference TdT synthesizes DNA. **(A.iii.)** This results in a change in the overall nucleotide incorporation preference upon a change in Ca^2+^. **(B) Top:** 100 μM CaCl_2_ was added to the extension reaction at 30 minutes to generate a 0→1 transition. **Center:** Mean output signal across 3 biological replicates. **Bottom:** Using a modified version of the algorithm detailed in Glaser et al. (13), the signal was deconvoluted into a binary response, with the predicted switch time of 30 minutes (**C) Top:** 50 μM EGTA was added to the extension reaction at 30 minutes to generate a 1→0 transition. **Center:** Mean output signal across 3 biological replicates. **Bottom:** Using a modified version of the algorithm detailed in Glaser et al. (13), the signal was deconvoluted into a binary response, with the predicted switch time of 22 minutes.

We employed the natural calcium sensing protein, Calmodulin (CaM), and a cognate binding peptide, M13, to generate a TdT with allosterically modulated activity. The calcium-dependent interaction between CaM and M13 has been previously utilized to engineer allosteric calcium biosensors (27, 28) and as a platform for generalizable ligand biosensors (29, 30).

Here, we generated variants with M13 fused to one of 4 sites in mTdT that were predicted to minimize the structural disruption of inserting the M13 sequence using SCHEMA-RASPP(31) (Fig S14, Table S2). After initial activity screening (Fig S15) of the variants we observed that one variant, mTdT(M13-388), retained polymerase activity. CaM was subsequently fused to the N-terminus of mTdT(M13-388) via linker. Primer extension reactions showed that the resulting CaM-mTdT(m13-388) variant was active in calcium-free conditions and inactive in calcium-added conditions (Fig S16). To confirm that the calcium-dependent interaction between CaM and M13 was responsible for the observed activity modulation, we mutated four essential Ca^2+^-binding residues in CaM, which ablated the calcium-sensitivity of CaM-mTdT(M13-388) (Fig S16).

Depending on the application, sensor polymerases with different calcium affinities may be useful to selectively record Ca^2+^ fluctuations exceeding threshold concentrations. We anticipated that the modular design of CaM-mTdT(M13-388) would allow the properties of the fusion to be rationally modified with CaM variants with known differences in Ca^2+^ affinity. Polymerase activity was determined by the length distributions of primer extensions, CaM-mTdT(M13-388) variants containing the CaM mutants D96V, D130G, and D142L which reduce the calcium affinity of CaM (32). We observed that all variants exhibited greater activity than the unmodified CaM-mTdT(m13-388) in the presence of low concentrations of calcium (Fig. S17). Moreover, the increase in activity correlated with the reported effective Ca^2+^ K_D_ of the variants, demonstrating that the calcium-sensitivity of CaM-mTdT(M13-388) can be rationally tuned.

Next, we tested if the TURTLES-2 system could the Ca^2+^ state into DNA. CaM-mTdT(M13-388) was purified and NGS analysis of extension reactions performed with the polymerase confirmed the calcium-sensitive phenotype (Fig. S18 & 19). We characterized CaM-mTdT(M13-388) in the context of the TURTLES-2 recording system by performing extensions with a mixture of purified bovine TdT and CaM-mTdT(M13-388) in calcium-free and calcium-added conditions. The two-polymerase system exhibited a significantly altered nucleotide selectivity in the calcium-added conditions (Fig S20). As expected, in the calcium-free condition the overall base incorporation preference was approximately the average of the observed preferences of bovine TdT and CaM-mTdT(M13-388) whereas the overall base preference in the calcium added condition was nearly identical to that of bovine TdT (Fig. S21). We conclude that the differential overall base selectivity of the TURTLES-2 system is capable of encoding the environmental calcium state into DNA.

### Recording a single step change in Ca^2+^ concentration with minutes resolution with two TdT system

We next investigated if the differential calcium response of TURTLES-2 could be used to infer the time at which calcium concentrations changed in an extension reaction. During a 60-minute extension we tested both 0→1 and 1→0 step transitions at 30 minutes, where ‘0’ is calcium-free and ‘1’ is calcium-added (Fig. 3B top, Fig. 3C top, Supplementary text 6). Using a variation of the model developed in Glaser et al. (13) we inferred transition times of 30 minutes for the 1→0 transition (Fig. 3B bottom) and 22 minutes for the 0→1 transition (Fig. 3C bottom). The decreased accuracy of the estimated time for the 0→1 transition did not correspond to an increase in measurement variability, suggesting that the offset is due to systematic factors. Time estimations for 0→1 transitions in TURTLES-2 would likely improve with more sophisticated decoding algorithms and deeper characterization of the transition behavior of CaM-mTdT(M13-388). We conclude that TURTLES-2 enables high resolution temporal encoding of calcium signals.

## Discussion

In this study, we demonstrated two DNA synthesis-based recording systems that encode and record the temporal dynamics of fluctuating environmental signals with minutes accuracy. By coupling sensing and writing functions, TURTLES simplifies the recording apparatus to post-translational system. This gives TURTLES distinct advantages over the temporal constraints of existing tools, enabling heretofore unprecedented temporal accuracy and resolution. While TURTLES-1 can record several physiologically relevant signals, TURTLES-2 lends tunability to the recording system with simple rational engineering. Given the uncomplicated and genetically-encodable design of TURTLES systems, we anticipate that they can be readily adapted for both *in vitro* and *in vivo* applications.

While many DNA-based biosensors have been deployed for studying physiological signals of interest (33–37), the scalability and spatial resolution of biosensors is intrinsically limiting in some applications (38). By leveraging *post hoc* recovery of biological data, optimized TURTLES systems may be capable of enabling otherwise inaccessible high-resolution spatial and temporal recordings of physiological signaling molecules that fluctuate on the timescale of 10^1^-10^3^ minutes. Such signals include slow calcium signaling that occurs in neurons (13, 14, 38, 39) and vertebrate development (22). Additional optimization of the TURTLES system will be required to enable the spatiotemporal resolution for characterizing systems with shorter timescales. Beyond biological applications, there has been a sustained interest in biosensors for testing environmental parameters such as water quality. For longer term tracking of contaminating metal ions in water, one could use TURTLES to track cobalt concentration over time (40, 41). In concert with microfluidic reaction control, TURTLES can also serve as a competitive platform for enzymatic DNA synthesis for data archiving (9–11), which is an appealing alternative to phosphoramidite methods due to the low cost and reduced environmental impact (42). Although TURTLES does not control the specific base added, as few as 5-10 bases could encode a bit. While this information density is lower than that of base-specific DNA synthesis but does not require specialized substrates or complex reaction cycling, which may yield cost savings with respect to base-specified methods.

Going forward, more sophisticated computational methods will improve the recording accuracy of TURTLES. In this study we utilized simple, intuitive models of TdT activity to transform sequence data into temporal information. By incorporating kinetic models of TdT activity or machine learning to classify signal changes along individual DNA strands, the accuracy of temporal estimations could likely be increased. These methods would also improve the robustness of TURTLES recordings by reducing the required sequencing depth from thousands to hundreds of reads. In this work, both the inputs and outputs for TURTLES were binarized, however the underlying principle can be extended to record continuously varying analog signals with improved decoding algorithms.

The quality of TURTLES recordings may also be improved by engineering the properties of TdTs. In particular, the sequencing depth required to accurately decode recordings can be reduced by increasing the magnitude of changes in nucleotide selectivity in response to inputs. Likewise, reference TdTs that have a more distinct nucleotide selectivity from the CaM-mTdT(M13-388) sensor TdT can be engineered or identified among natural TdT diversity. Improvements to the temporal resolution of TURTLES systems can be accomplished by enhancing the nucleotide incorporation rate of TdT. In TURTLES-2, structural optimization of CaM-mTdT(M13-388) may improve temporal resolution by optimizing the kinetics of the CaM-M13 interaction. Notably, fluorescent biosensors based on CaM-M13 interactions can report calcium spikes on the order of seconds(43, 44), suggesting that calcium sensing will not be limiting with respect to temporal resolution in an optimized system. The functionality of TURTLES-2 may be further expanded by employing generalizable sensors based on the CaM-M13 interaction (29), or by probing TdT with sensing domains other than calmodulin such that new signals of interest can be encoded or recorded. In all, we have demonstrated a new methodology for recording dynamic, environmental information into DNA that relies only on allosteric regulation, enabling minutes resolution.

## Materials and Methods

### Enzymes and ssDNA substrate

Terminal deoxynucleotidyl polymerase, T4 RNA ligase I, Phusion High-Fidelity PCR Master Mix with HF Buffer were purchased through New England Biolabs (NEB). ssDNA substrates used for extension reactions were ordered from Integrated DNA Technologies (IDT) with standard desalting. dNTPs were obtained from Bioline.

### CaM fusion design and screening

Four fusion proteins were designed that consisted of CaM fused to the N terminus of mTdT by a (GGGGS)_4_ linker and M13 inserted immediately following the fusion residue (see below) with flanking GS linkers. Fusion sites were selected from crossover sites identified with the SCHEMA/RASPP algorithm based on which sites were in catalytically essential regions and would be sterically available to CaM. SCHEMA crossover sites were calculated according to previously described protocols(31). Sets of crossover points were calculated for 3, 4, 5, 6, and 7 total crossovers. Calculations were performed with the following parameters: minimum fragment length = 4, bin width = 1, parent sequence = NP_001036693.1, parent structure = PDB 4i27 (all ligands, metals, and waters removed), homology sequences = NP_803461.1, AAH12920.1, NP_001012479.1, [XP_021064401.1](https://www.ncbi.nlm.nih.gov/protein/NP_001012479.1?report=genbank&log$=protalign&blast_rank=10&RID=R9BB76AK014), XP_020136193.1. All sequences were trimmed to only include residues crystallized in the parent structure. Fusion sites were selected from crossover points that were in the DNA-binding region of mTdT (residues 282, 284, 287) or in Loop 1, a catalytically essential structure (residue 388). M13 fusions were screened for activity without N-terminal CaM to validate that the fusion was tolerated.

### Cloning CaM-TdT(M13) variants

Molecular cloning of DNA constructs was completed under a contract research agreement with the lab of Dr. J. Andrew Jones at Miami University – Oxford, OH. The pET28a-M-CaM-cTdTS-M13-XXX (282, 284, 287, and 388) variants were constructed using a two-part Gibson Assembly method. The approximately 75bp M13 fragment was amplified from linear double stranded DNA template (gBlock-CaM-Linker-M13, IDT) using Accuzyme DNA polymerase (Bioline) using DNA primers P21 – P28 listed in Table 1. The amplicon was then purified using a Cycle Pure Kit (Omega Biotek). The vector backbone fragment was amplified from pET28a-M-CaM-cTdTS plasmid DNA constructed above using *PfuUltra* II Hotstart PCR Master Mix using DNA primers P29 – P36 listed in Table 1. The PCR product was then digested with *Dpn*I to remove DNA template. The approximately 8100bp amplicon was purified using a gel extraction kit (Omega Biotek). DNA concentration of both linear fragments was measured using the Take3 plate coupled with a Biotek Cytation 5 plate reader. Corresponding backbone and M13 fragments were then assembled using the repliQa™ HiFi Assembly Kit (Quanta bio), transformed into chemically competent DH5α, and selected on LB-Kanamycin (50 μg/mL) plates. Individual colonies were then screened via restriction digestion and verified using Sanger sequencing (CBFG – Miami University) with primers S1 – S8, Table 1.

### CaM-TdT(M13-388) expression and purification

Purification optimizations determined that N-terminal MBP was unnecessary for expression and purification and was not included in the final expression construct. The expression construct (pET28a-CaM-mTdT(m13-388)) was transformed into chemically competent NEB T7Express cells, plated on kanamycin selective plates, and incubated at 37°C. The following day, a single colony was selected and inoculated into 5mL of kanamycin supplemented LB. the culture was incubated for 20 hours at 37°C. Four flasks of 120mL kanamycin supplemented LB were inoculated 1:400 (v/v) with the overnight culture. The cultures were incubated with shaking at 250 RPM. Once the OD600 was between 0.5 and 0.6, the cultures were cooled to room temperature and induced with 1mM IPTG. Following induction, the cultures were incubated for 18 hours at 15°C. The cells were pelleted at 4°C and the supernatant was discarded. The decanted cell pellets were stored at -80°C.

The cell pellets were thawed on ice. Lysis and affinity chromatography were performed using the Takara Bio HisTALON gravity column purification kit; all steps were performed according to the manufacturer’s native protein extraction protocol. Note that the cell pellets were treated with optional DnaseI and lysozyme during lysis. 1mL of Takara Bio TALON metal affinity resin was used for affinity chromatography. All binding and wash steps were performed on ice with shaking at 250 RPM. 15 bed volumes of wash buffer were used for all washes. CaM-mTdT(M13-388) was eluted from the resin in 10, 500uL fractions. Each fraction was analyzed by SDS-PAGE, and the total protein concentration in each fraction was measured by absorbance at 280nm. The first five elution fractions, which contained the majority of eluted protein, were pooled.

The pooled fractions were diluted in binding buffer (20 mM Tris-HCl, pH 8.3) and further purified by anion exchange chromatography using a Cytiva HiTrap Q HP 5mL column and a 40 CV gradient from 0 mM to 1 mM NaCl in binding buffer with a GE Healthcare AKTAxpress FPLC. The protein eluted in two fractions.

Both elutions were buffer exchanged by dialysis into a storage buffer consisting of 200mM KH_2_PO_4_ and 100mM NaCl at pH 6.5 and concentrated using Vivaspin 20 columns to a final concentration of 0.37 mg/mL for the first fraction and 0.98 mg/mL for the second fraction. The fractions were aliquoted, and flash frozen on dry ice for storage at -80°C. Notably, PAGE analysis showed that the second elution contained a product at 25kDa in addition to the expected fusion protein at approximately 70kDa. Both CaM-mTdT(M13-388) elutions recapitulated the calcium-sensitive phenotype and exhibited similar nucleotide selectivity (Fig S19-20). As significantly more protein was recovered in the second elution it was used for all subsequent experiments.

### Cell free protein expression and primer extension assay

For initial activity screening of fusion variants and CaM-mTdT(M13-388) characterization, proteins were expressed in cell-free reactions. Variants were expressed using NEB PURExpress in 25 μL reactions containing 40% (v/v) PURExpress Solution A, 30% (v/v) PURExpress Solution B, 1.6 U/μL NEB Rnase I, 10 ng/μL expression vector DNA, and dH_2_O to volume. The expression reactions were incubated for 4 hours at 30°C.

Primer extension reactions were prepared on ice. Primer extensions were performed in 25 μL reactions containing 1X NEB TdT Reaction buffer, 0.8 μM single-stranded, FAM-labelled substrate DNA FAM_NB (Table S2), 1mM dNTPs, polymerase, and dH_2_O to 25 μL. For variants expressed in PUREexpress, 2.5 μL of the expression reaction was used immediately after expression; 20U (approximately 0.2 μg) of the NEB bovine TdTwas used for positive control reactions; approximately 0.5 μg, of purified CaM-mTdT(M13-388) was used for activity validation reactions after purification. For calcium-added conditions, CaCl_2_ was added to the reactions to a final concentration of 1mM. Extension reactions were incubated for 2 hours at 37°C

Completed extensions were analyzed by urea-PAGE. 8 μL of each completed primer extension reaction was combined with 12 μL of BioRad 2x TBE urea sample buffer and boiled for 10 minutes. The boiled samples were loaded onto a 10% polyacrylamide TBE urea gel (bioRad 4566036), and 200V was applied to the gel for 40 minutes. The gels were imaged on a GE Healthcare Typhoon 9400 laser scanner using a 200um pixel size and λ_ex_=488nm, λ_em_=520nm BP40. Imaging gain was adjusted for each experiment to avoid saturation.

### Extension reaction for calculating effect of Co^2+^, Ca^2+^, Zn^2+^, and temperature on overall dNTP preference of TdT

Each extension reaction consisted of a final concentration of 10 μM ssDNA substrate CS1 (Table S2), 1 mM dNTP mix (each dNTP at 1 mM final concentration), 1.4x NEB TdT reaction buffer, and 10 units of TdT to a final volume of 50 μL. When testing the effect of cations, CoCl_2_ was added at a final concentration of 0.25 mM, CaCl_2_ at 2 mM, or Zn(Ac)_2_ at 20 μM. It is important to note that reaction initiation was done by adding TdT to the ssDNA substrate mix (ssDNA substrate mix consisted of the ssDNA substrate, dNTPs and the cation). Prior to reaction initiation, the ssDNA substrate mix and TdT were stored in separate PCR strip tubes at 0 °C (on ice). The reaction was run for 1 hour at 37 °C in a Bio-Rad PCR block. When testing the effect of temperature, the same reaction mix was run on a Bio-Rad PCR block set at tested temperatures for 1 hour. Reactions were stopped by freezing at -20 °C. For initial testing, reactions were analyzed by urea-PAGE 2 μL of the reaction was mixed with 12 μL of TBE-Urea (Bio-Rad) loading dye and boiled for 10 minutes at 100 °C. All of the diluted extension reaction was then loaded onto 30 μL, 10 well 10% TBE-Urea Gel (Bio-Rad) and run for 40 minutes at 200 V. Immediately after the run was over, the gel was stained with Sybr Gold for 15 minutes and imaged on an ImageQuant BioRad.

### TURTLES 0→1 extension reactions

Mg^2+^ only for 1 hour (signal 0) and Mg^2+^+Co^2+^ for 1 hour (signal 1) were set up as regular extension reaction mentioned above. The 0→1 reactions where the signal changed from 0 to 1 at various times during the 1 hour extension were run starting at a total volume of 45 μL with Mg^2+^ only. 5 μL 2.5 mM CoCl_2_ was added at the time we wanted the signal to change from 0 to 1. Reactions were all run for a total of 1 hour in triplicates. Fresh signal 0 and signal 1 controls were run with each set-up.

### TURTLES-2 controls and 0→1 and 1→0 extension reactions

TURTLES-2 extension reactions contained 1X NEB TdT reaction buffer, 0.1 μM ssDNA substrate CS1_5N (Table S2), 1 mM dNTP mixture, and 2.5 μL polymerase mixture. The polymerase mixture contained CaM-mTdT(M13-388) at a concentration of 0.45 mg/mL and NEB TdT at a concentration of 0.002 mg/mL (0.4U). Calcium-free reactions included EGTA at a final concentration of 50 μM. The high calcium control for 0→1 reactions was supplemented with CaCl_2_ and EGTA to final concentrations of 100 μM and 50 μM, respectively. The high calcium control for 1→0 reactions was neither supplemented with EGTA nor CaCl_2_ (supplementary text 6). Reactions were brought to a final volume of 25 μL with nuclease-free water. Reactions were assembled on ice and initiated by adding TdT to the substrate mixture. Reactions were incubated for 1 hour at 37 °C in a Bio-Rad PCR block and terminated by heating reactions to 80 °C for 10 minutes.

Signal transitions were performed by 1uL additions at 30 minutes. for 1→0 reactions, the addition contained 1X NEB TdT buffer and 1.3 mM EGTA (50 mM EGTA in final reaction post- addition). for 1→0 reactions, the addition contained 1X NEB TdT buffer and 2.6 mM CaCl_2_ (100 μM CaCl_2_ in final reaction post- addition).

### Extension reactions for 0→1→0 set-up

Mg^2+^ only for 1 hour (signal 0) and Mg^2+^+Co^2+^ for 1 hour (signal 1) were set up as regular extension reaction mentioned above. The 0→1→0 reactions where the signal changed from 0 to 1 at 20 minutes and back to 0 at 40 minutes were run starting at a total volume of 45 μL with Mg^2+^ only. 5 μL 2.5 mM CoCl_2_ was added at the time we wanted the signal to change from 0→1. For changing the signal from 1→0, since the ssDNA was suspended in reaction buffer for these set-ups, we used a ssDNA clean up kit (methods mentioned below) to remove the reaction buffer, TdT, cation and dNTPs from each reaction. All of the ssDNA collected from the ssDNA clean up kit (20 μL) was then prepared for the last part of the extension reaction. Collected ssDNA was mixed with a dNTP mix at a final concentration of 1 mM (each dNTP at 1 mM final concentration), 1.4x TdT reaction buffer and 10 units of TdT to a final volume of 50 μL. All reactions were always initiated by adding TdT in the end. Signal 0 and signal 1 controls were run for 1 hour for each set-up in triplicates and also put through the ssDNA wash step at 40 minutes. Six replicates were run for 0→1→0 reactions.

### ssDNA wash for replacing buffers for 0→1→0 reactions

For changing cation concentration from 1 to 0 we utilized the ssDNA clean-up kit (ssDNA/RNA clean/concentrator D7010) from Zymo Research such that all the extended ssDNA synthesized in the initial part of the experiment was retained on the column and the TdT, reaction buffer, cation and dNTPs were washed away. Each 50 μL extension reaction was individually loaded into a separate column. Protocol was followed as mentioned in the kit. ssDNA was eluted into 20 μL ddH_2_O. We noticed in initial tests that after using the ssDNA clean-up kit, there was little to no TdT-based extension in some replicates (data not included). We presume this is due some ethanol getting carried forward into the eluted ssDNA. Thus we extended the dry spin time based on suggestion from Zymo Research to 4 minutes. We also utilized two other ways to evaporate any remaining ethanol after the column dry spin step based on protocol mentioned in Cold Spring Harbor Protocols(45). We either kept the columns open in a biohood for 15 minutes to allow for evaporation, or after elution of ssDNA we kept the 1.5 mL eppendorf tubes containing the eluted ssDNA open at 45 °C for 3 minutes. Both methods gave better ethanol removal than just dry spin, and they were tried in triplicates and averaged and plotted for the time prediction analysis (Fig. 3C).

### Illumina library preparation and sequencing

Our sample preparation pipeline for NGS was adapted from a previous protocol(46),(47). After extension reaction, 2 μL of the product was utilized for a ligation reaction. 22 bp universal tag, common sequence 2 (CS2) of the Fluidigm Access Array Barcode Library for Illumina Sequencers (Fluidigm), synthesized as ssDNA with a 5’ phosphate modification and PAGE purified (Integrated DNA Technologies), was blunt-end ligated to the 3’ end of extended products using T4 RNA ligase. Ligation reactions were carried out in 20 μL volumes and consisted of 2 μL of extension reaction, 1 uM CS1 ssDNA, 1X T4 RNA Ligase Reaction Buffer (NEB), and 10 units of T4 RNA Ligase 1 (NEB). Ligation reactions were incubated at 25 °C for 16 hours. Ligated products were stored at −20 °C until PCR that was carried out on the same day. Ligation products were never stored at -20°C for more than 24 hours.

PCR was performed with barcoded primer sets from the Access Array Barcode Library for Illumina Sequencers (Fluidigm) to label extension products from up to 96 individual reactions. Each PCR primer set contained a unique barcode in the reverse primer. From 5’-3’ the forward PCR primer (PE1 CS1) contained a 25-base paired-end Illumina adapter 1 sequence followed by CS1. The binding target of the forward PCR primer was the reverse complement of the CS1 tag that was used as the starting DNA substrate. From 5’-3’ the reverse PCR primer (PE2 BC CS2) consisted of a 24-base paired-end Illumina adapter 2 sequence (PE2), a 10-base Fluidigm barcode (BC), and the reverse complement of CS2. CS2 DNA that had been ligated onto the 3’ end of extended products served as the reverse PCR primer-binding site. Each PCR reaction consisted of 2 μL of ligation product, 1X Phusion High-Fidelity PCR Master Mix with HF Buffer (NEB), and 400 nM forward and reverse Fluidigm PCR primers in a 20 μL reaction volume. Products were initially denatured for 30 s at 98 °C, followed by 20 cycles of 10 s at 98 °C (denaturation), 30 s at 60 °C (annealing), and 30 s at 72 °C (extension). Final extensions were performed at 72 °C for 10 min. Amplified products were stored at −20 °C until clean up and pooling. QC for individual sequencing libraries was performed as follows. 2 μL of each library was pooled into a QC pool and the size and approximate concentration was determined using Agilent 4200 Tapestation. Pool concentration was further determined using Qubit and qPCR methods. Sequencing was performed on an Illumina MiniSeq Mid Output flow cell and sequencing was initiated using custom sequencing primers targeting the CS1 and CS2 conserved sites in the library linkers. Additionally phiX control library was spiked into the run at 15-20% to increase diversity of the library clustering across the flow cell.

After demultiplexing, the percent seen for each sample was used to calculate a new volume to pool for a final sequencing run with evenly balanced indexing across all samples. This pool was sequenced with metrics identical to the QC pool. Library preparation and sequencing were performed at the University of Illinois at Chicago Sequencing Core (UICSQC).

### NGS Data Preprocessing

For each sample, the NGS reads were first trimmed and filtered using cutadapt (v1.16). Only NGS read pairs with both Illumina Common Sequence adapters, CS1 and CS2, were kept. Of these, CS2 was trimmed off each R1 sequence and CS1 was trimmed off each R2 sequence. Cutadapt parameters were set as following: a minimum quality cutoff (-q) of 30, a maximum error rate (-e) of 0.05, a minimum overlap (-O) of 10, and a minimum extension length (-m) of 1. The minimum overlap was set to be higher than the default value of 3 because extended sequences in this case are random, and we did not want to filter out sequences where the final 1-10 bases just happen to look like the first 10 bases of CS2 (the read must still contain a full CS2 sequence for it to be kept and subsequently trimmed, however). The 3’ (-a) adapter trimmed from the R1 reads was 5’AGACCAAGTCTCTGCTACCGTA3’ (CS2 reverse complement), and the 5’ (-A) adapter trimmed from the R2 reads was 5’TGTAGAACCATGTCGTCAGTGT3’ (CS1 reverse complement). FastQC was used to quickly inspect the output trimmed .fastq files before downstream analysis. See *filter_and_trim_TdT.sh* at https://github.com/tyo-nu/turtles for an example preprocessing script. All runs were trimmed using this script. All initial preprocessing was done on Quest, Northwestern University’s high-performance computing facility, using a node running Red Hat Enterprise Linux Server release 7.5 (Maipo) with 4 cores and 4 GB of RAM, although only 1 core was used. Preprocessing took between 5 and 30 minutes depending on the number of conditions, replicates, and reads per replicate in a given run.

Finally, for each analysis, we did further preprocessing locally. We cut off bases that were still present in the reads but not added during the experiment. Degenerate bases (if any) that are part of the 5’ ssDNA substrate (at its 3’ end before the extension) were removed from the beginning of each sequence. Then, we cut off 5.8 bases off the end of every sequence because we found that, on average, 5.8 bases were being added after the extension reaction during the 16 hour ligation step (Figure S14). Because 5.8 is not an integer value, we cut 5 bases off of 80% of the sequences and 6 bases off of 20% of the sequences. We also filtered out sequences with length less than 6 bases.

### Timepoint prediction for 0→1 single step change experiment

All further analysis was done in python using Jupyter Notebooks. You can find all the Jupyter Notebooks used for this publication at https://github.com/tyo-nu/turtles. The following algorithm was applied in order to (1) read and normalize each sequence by its own length, (2) calculate a distance metric using the relative dATP, dCTP, dGTP, and dTTP percent incorporation changes between each condition and the 0 control, and (3) transform distances for all conditions into 0 → 1 space based on the 0 and 1 control distance values.

We first normalize each sequence by length, such that all bases in each sequence are counted across 1000 bins. For example, for a sequence of length 10, the first base would get counted in the first 100 bins, the next base in bins 100-200, and so on.

We then calculate base composition, *X*_*ij*_, in the sequence for condition, *i*, at each bin with position, *j*, using the formula for a closure (equation 1). Note that *i* is unique for each (condition, replicate) pair if multiple replicates are present for a given experimental condition.

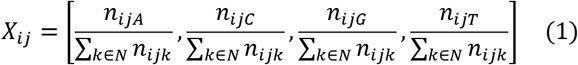

Here, *n*_*ijk*_ is the total count of dATP, dCTP, dGTP, or dTTP depending on the value of *k* (*k* ∈ *N* = {*A*, *C*, *G*, *T*}) across all sequences for condition, *i*, at bin, *j*.

To calculate distance between two compositions at a given bin location (e.g. between the 0 and 1 controls at every bin), we have to first transform the compositional data. We cannot simply take the L2 norm difference of each compositional element because the elements of a composition violate the principle of normality due to the total sum rule (all elements add up to 100%). Thus, the data is first transformed by using the center log-ratio (clr) transformation which maps this 4-component composition from a 3-dimensional space to a 4-dimensional space. We then take the L2 norm of these transformed normal elements. This distance metric is known as the Aitchison Distance, which is used here to calculate the base composition distance, *d*_*j*_(0, *i*), from the 0 control to each condition, *i*, at each bin, *j* (equation 2).

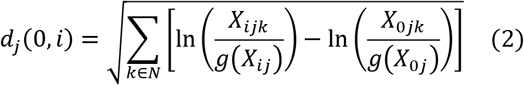

*N* = {*A*, *C*, *G*, *T*} and *g*(*X*_*ij*_) is the geometric mean for condition, *i*, and bin, *j*, across all four bases in *N* (equation 3).

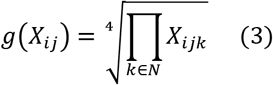

For condition, *i*, and bin *j*, the output signal, *s*_*ij*_, is calculated as

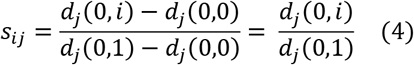

where *d*_*j*_(0,1) is the Aitchison distance between the 0 control base composition and 1 control base composition at bin, *j*. *d*_*j*_(0,0) = 0 for all *j*. If there were multiple replicates for the 0 control, their average composition was used for *X*_0*j*_ (and *X*_0*jk*_) in equation 2. If there were multiple replicates for the 1 control, their average composition was similarly used to calculate *d*_*j*_(0,1) in equation 4.

Next, the switch times were estimated for each condition, *i*, which contains a change in output signal, *s*_*ij*_, (e.g. via addition of Co halfway through the reaction). For experiments with more than one change (e.g. 0 → 1 → 0), a more sophisticated approach was used and is detailed below. However, the following simpler, more intuitive approach was used to predict switch times for 0 → 1 and 1 → 0.

Switch times were estimated for a given condition, *i*, by (1) finding 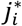, the average location across all the sequences (bin position, *j*) at which half the 1 control output signal is reached (i.e. *s*_*ij*_ = 0.5), (2) calculating *α*, the ratio of the average rate of nucleotide addition for the 0 and 1 controls, and (3) using 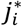 and *α* to calculate the switch time, 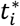, using equations 5 and 6. For a derivation of equation 5, see supplementary methods.

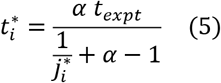

where

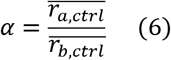

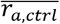 is the average synthesis rate of the first environmental condition before the switch. For example, 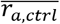 would be calculated using the 0 control for the condition, 0 → 1, but the 1 control for the condition, 1 → 0. The average synthesis rate is calculated by dividing the average extension length by the duration of the experiment. 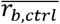 is the average synthesis rate for the second environmental condition (after the switch).

### Timepoint estimation for 0→1→0 multiple fluctuations experiment

To predict the Co^2+^ condition in the 0→1→0 experiment, we used the algorithm we developed in Glaser et al. for decoding continuous concentrations(13). The input to this algorithm is the amount of output signal on every nucleotide. Here, the output signal is *s*_*ij*_ from the previous section. The algorithm uses this information to predict continuous values of Co^2+^ between 0 and 1 for all time points that are most likely to produce the amount of output signal on the nucleotides. To binarize these predictions, we then set a threshold of 0.5. To be able to predict the values of Co^2+^, the algorithm requires knowledge of the expected amount of output signal in the 0 and 1 control conditions. Here, this is the average output signal across nucleotides in the 0 or 1 control experiments. The algorithm also requires knowledge of the rate of nucleotide addition. Here, we fit an inverse Gaussian distribution to the average experimental dNTP addition rate distribution (the distribution of the sequence lengths divided by the experiment time) from the control experiments. Note that this algorithm also assumes that the rate of dNTP addition is independent of the cation concentration. Thus, when making predictions in the 0→1→0 experiment, we do not account for differences in the rate of dNTP addition distributions between the 0 and 1 conditions. A future algorithm that takes this difference into account could yield more accurate predictions.

### *In silico* simulations of recording faster and higher number of input signal changes

Using the average dNTP incorporation rate from experiments, and the amount of output signal in the control conditions, we simulated additional experiments *in silico*. Each simulated experiment had at least 6 signal changes (instances of a single signal change from 0→1 or 1→0), where each condition was randomly chosen to be 0 or 1. All nucleotides that were added during the 0 or 1 condition had the signal associated with these control conditions. More specifically, to account for the experimental variability in signals within a given control condition, nucleotide signals were sampled from a Normal distribution determined by the experimental variability of nucleotide signals within the control conditions. We calculated the variability in two ways, corresponding to the two representative curves in Fig. S13A and S13C. In one, the variability was calculated across the first 100 nucleotides, in which there were at least 2000 recordings of all base numbers. In the second, the variability was calculated across the first 50 nucleotides, in which there were at least 60000 recordings of all base numbers. Using the output signal of the simulated nucleotides, we used the algorithm we developed in Glaser et al. for decoding binary concentrations (13). Accuracy corresponds to the percentage of conditions correctly classified as 0 or 1 over the duration of the entire recording experiment.

### Time point estimation for 0→1, 1→0 single step change for TURTLES-2 using an inverse model

To predict the Co^2+^ condition in the 0→1→0 experiment, we used a variation of the algorithm we developed in Glaser et al. for decoding continuous concentrations (13). This algorithm will predict continuous values of Co^2+^ between 0 and 1 for all time points that are most likely to produce the amount of signal. Here, instead of using the amount of signal on every nucleotide to predict the continuous concentrations, we use the normalized signal.

Let *s*_*ij*_ be the signal as a function of condition, *i*, and normalized position, *j*. Let *γ*_*ij*_(*t*) be the probability that a nucleotide corresponding to normalized position *j* was written at time *t*. Let ***C***_***i***_ be the normalized cation concentration for condition *i*. Like in (13), our model is that 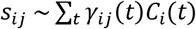. We use maximum likelihood estimation to find ***C***_***i***_ that minimizes 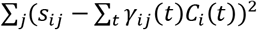 subject to *C*_*i*_(*t*) ∈ [0,1] and given condition, *i*. To binarize the predictions, we then set a threshold for ***C*** of 0.5.

Here, for an experiment of duration *t*_*expt*_ (e.g. 60 minutes), we let *γ*_*ij*_ = Ν((*j* − 0.5)/*t*_*expt*_, *σ*)/*Z*, where *Z* renormalizes the probability distribution after values outside the domain of [0, *t*_*expt*_] are set to 0. We set *σ* for each experiment so that *γ*_*i*1_(*t*_*expt*_) is equal to the frequency of strands with a single nucleotide divided by *t*_*expt*_ (because a normalized position of 1 would generally only be written in the *t*_*expt*_^th^ minute when there is a single nucleotide strand). Note that future work that more accurately models the kinematics of the polymerase to get a more accurate estimate of *γ* will provide improved results.

When running this algorithm on the Ca^2+^ data which has different rates when Ca^2+^ is or isn’t present, following the prediction of Ca^2+^ over time with the above algorithm, we used the ratio of incorporation rates between the 0 and 1 condition, as described by Equation 5 and 6, to rescale the results.

## Data and code availability

All data generated during this study are included in this published article and its Supplementary Information. NGS data are available from Sequence Read Archive https://www.ncbi.nlm.nih.gov/sra/PRJNA542184

## Acknowledgements

The authors would like to acknowledge Marija Milisavljevic for help with some experiments, and Bradley Biggs for helpful discussions and comments on the manuscript. This research was supported in part through the computational resources and staff contributions provided for the Quest high performance computing facility at Northwestern University, which is jointly supported by the Office of the Provost, the Office for Research, and Northwestern University Information Technology. All next generation sequencing was done with the help of the Next Generation Sequencing Core facility at University of Illinois at Chicago. Sanger sequencing was supported by the Northwestern University NUSeq Core Facility. Gel imaging was supported by the Northwestern University Keck Biophysics Facility and a Cancer Center Support Grant (NCI CA060553). The Keck Biophysics Facility’s Azure Sapphire Imager was funded by the 1S10OD026963-01 NIH grant. Protein purification was supported by the Northwestern University Recombinant Protein Production Core. This work was funded by the National Institutes of Health grants R01MH103910 (to KEJT, KPK, ESB and GMC), and UF1NS107697 (to KEJT, KPK, ESB) and National Institutes of Health Training Grant (T32GM008449) through Northwestern University’s Biotechnology Training Program (to JS and AC).

## Supplementary Information Text

**Supplementary Methods**: Extended description of methods

### 1. Extension reaction with individual dNTPs for testing effect of Co^2+^

For initial testing to show Co^2+^ dependent dNTP preference change the ssDNA substrate used was AMD006 (Table S2). Total reaction volume was 25 μL with 0.1 μM ssDNA substrate, 1x NEB TdT reaction buffer, and 0.1 mM of each dNTP tested. Final concentration of CoCl_2_ in the test reaction was 0.25 mM. Reactions were initiated by addition of 5 units of TdT per reaction. Reactions were run for 30 minutes at 37 °C and stopped by boiling at 70 °C for 10 minutes. Then, 8 μL of the reaction was mixed with 12 μL of TBE-Urea loading dye and boiled for 10 minutes at 100 °C. All of the diluted extension reaction was then loaded onto 30 μL, 10 well 10% TBE-Urea Gel (Bio-Rad) and run for 40 minutes at 200 V. Immediately after the run was over, the gel was stained with Sybr Gold for 15 minutes and imaged on ImageQuant BioRad.

### 2. Extension reactions for 1→0 set-up

Mg^2+^ only for 1 hour (signal 0) and Mg^2+^+Co^2+^ for 1 hour (signal 1) were set-up as regular extension reactions mentioned in Materials and Methods. The 1→0 reactions where the signal changed from 1 to 0 at 40 minutes were put through a ssDNA was step at 40 minutes. ssDNA wash to remove cations, TdT and dNTPs was done exactly as mentioned in Materials and Methods. Reactions were all run for 1 hour in triplicates. Signal 0 and signal 1 controls were run for 1 hour for each set-up in triplicates and also put through the ssDNA wash step at 40 minutes.

### 3. Derivation of Equation 5

We start by deriving the equations for the average rate before the switch (*r*_*A*_) and after the switch (*r*_*B*_) for condition, *i*:

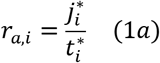

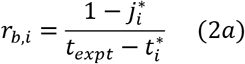

where 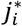 is the average location in the sequences (length fraction, 0 to 1) at which the output signal, *s*_*ij*_, reaches 0.5 (Equation 4), 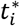 is the switch time, and *t*_*expt*_ is the total duration of the experiment. Because we can estimate *r*_*a*,*i*_ and *r*_*b*,*i*_ from average rates of the 0 and 1 controls across replicates (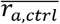 and 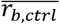), we can use their ratio to combine equation 1a and 2a, above to write

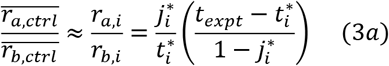

Solving for 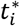, we get equation 5:

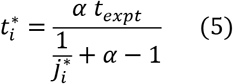

where

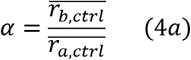

We use equation 5 for time prediction (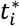) after calculating 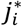 for a given condition and *α* from the 0 and 1 controls. In equation 4a, *a* is the first condition before the switch (0 or 1) and *b* is the condition after the switch (1 or 0).

### 4. Extensions reaction set-up for calculating rate of dNTP addition

Each extension reaction consisted of a final concentration of 10 μM initiating ssDNA substrate, 1 mM dNTP mix (each dNTP at 1 mM final concentration), 1.4x NEB TdT reaction buffer, and 10 units of TdT to a final volume of 50 μL. The ssDNA substrate used for this extension reaction was CS1_5N. We have shown (data not included) that the identity of the last 5 bases on the 3’ end of the substrate affects the identity of the dNTP added to the ssDNA substrate. Thus, we purchased a ssDNA substrate (CS1_5N) with the last 5 bases having the base composition same as TdT dNTP preference under signal 0 (25% dATP, 15% dCTP, 45% dGTP and 15% dTTP). The reactions were initiated upon addition of TdT and run at 37 °C for 2 hours. 2 μL of sample was collected and immediately frozen (on ice, 0 °C) at 30 s, 1 min, 2 min, 3 min, 4 min, 5 min, 10 min, 20 min, 30 min, 45 min, 60 min, 92 min and 120 min. Subsequently, each sample was put through the ligation and Illumina library generation process as mentioned in Materials and Methods.

### 5. Test set-up for checking ssDNA clean-up kit bias

Mg^2+^ only for 1 hour (signal 0) and Mg^2+^+Co^2+^ for 1 hour (signal 1) were set up as regular extension reactions mentioned in Materials and Methods. The 0→1 reactions where the signal changed from 0 to 1 during the 1 hour extension were run starting with 45 μL with Mg^2+^ only. 5 μL of 2.5 mM CoCl_2_ was added at 10 min. Reactions were all run for 1 hour in triplicates. Fresh signal 0 and signal 1 controls were run for 1 hour with each set-up. 2 μL of extension reaction was used for ligation (“No Wash” set of samples). Ligation and subsequent PCR steps for Illumina library generation were followed as mentioned in Materials and Methods. Rest of the 48 uL of extension reaction was washed using the ssDNA clean-up kit. Protocol was followed as mentioned in the kit. ssDNA was eluted into 25 μL of ddH_2_O and 2 μL of that was used for ligation (“Wash” set of samples). Ligation and subsequent PCR steps for Illumina library generation were followed as mentioned in Materials and Methods. Data obtained from Illumina sequencing was analyzed for the “No Wash” and “Wash” set of samples. Further, switch time calculations were carried out as mentioned previously (Fig. S12).

### 6. High calcium conditions for TURTLES-2 reactions

High calcium conditions for 0→1 and 1→0 reactions had different compositions to enable transitions without employing intermediate column washes. Although commercial TdT reaction buffer contains no added calcium, we observed that CaM-mTdT(M13-388) was inactivated in reactions that were not supplemented with at least 50 μM EGTA (Figure S18), very likely due to calcium present in water or other reagents used. By titrating EGTA into a TdT extension reaction supplemented with 7 μM fura-2 until the fura-2 signal plateaued, we estimated that most free calcium could be eliminated from the reactions by the addition of 50 μM EGTA. By employing the un-supplemented reaction buffer as the high calcium condition for 1→0 reactions, we were able to transition to a low calcium condition with the addition of 50 μM EGTA.

