## Supplementary Figures and Tables for "Recording Temporal Signals with Minutes Resolution Using Enzymatic DNA Synthesis"

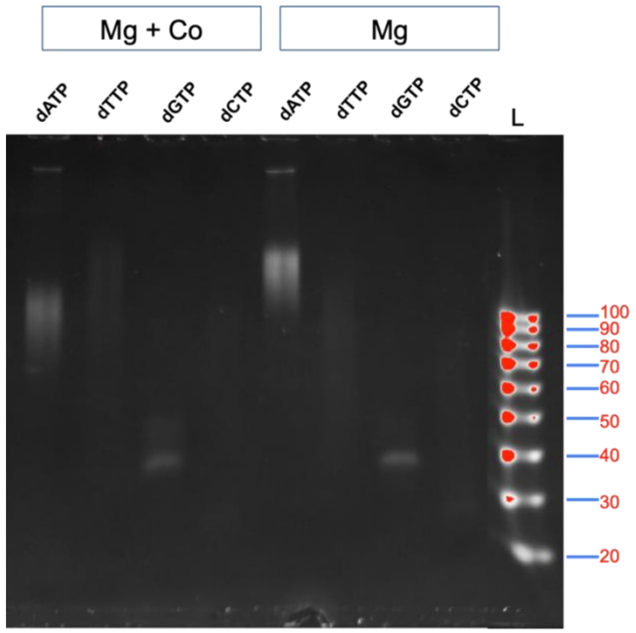


**Figure S1: Testing change in individual dNTP preference upon $Co^{2+}$ addition**
ssDNA substrate extensions carried out by TdT using just dATP, dTTP, dGTP, or dCTP in presence of $Mg^{2+}$+$Co^{2+}$ (first 4 lanes) or in presence of just $Mg^{2+}$ (next 4 lanes) were run on a gel. “L” is ssDNA size marker. Reactions were carried out as mentioned in supplementary text.

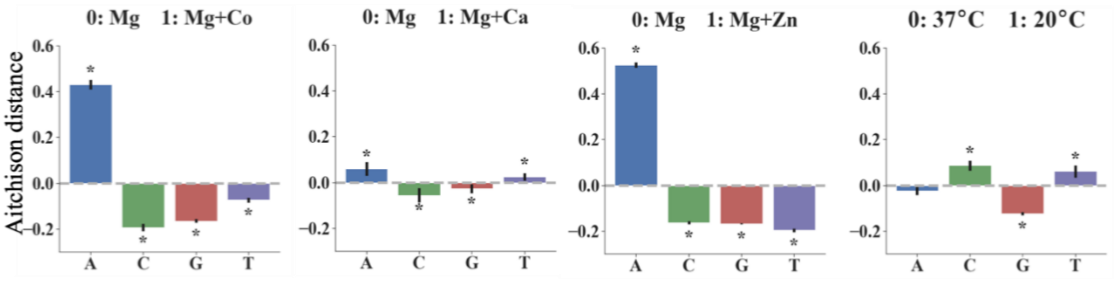


**Figure S2: Aitchison distance for dATP, dCTP, dGTP and dTTP incorporation by TdT in the presence or absence of various signals.** Signal 0 is always 10 mM $Mg^{2+}$ at 37 °C for 1 hour. Signal 1 was, going from left to right: (1) 10 mM $Mg^{2+}$ + 0.25 mM $Co^{2+}$ at 37 °C for 1 hour; (2) 10 mM $Mg^{2+}$ + 1 mM $Ca^{2+}$ at 37 °C for 1 hour; (3) 10 mM Mg + 20 μM $Zn^{2+}$ at 37 °C for 1 hour; and (4) 10 mM $Mg^{2+}$ at 20 °C for 1 hour. Error bars show two standard deviations of the mean Aitchison distance. Statistical significance was assessed after first transforming the data into Aitchison space which makes each dNTP frequency change statistically independent of the others. All base incorporation changes were found to be statistically significant, as shown by asterisks, ($\alpha = 0.01$) except for the change in dATP for 20 °C signal (p-value = 0.019).

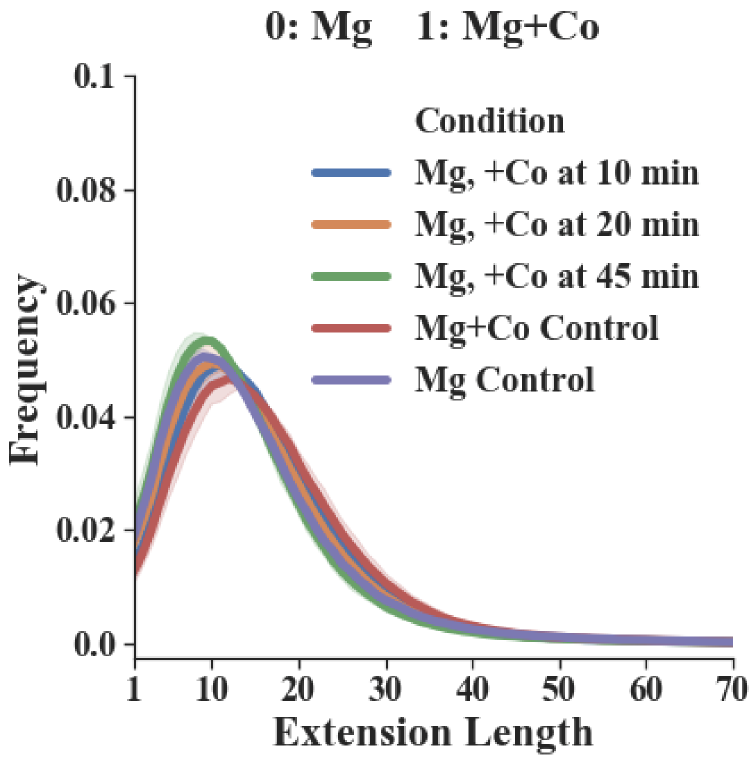


**Figure S3: Length distribution of extensions upon addition of $Co^{2+}$ based on NGS data.**
We calculated the mean frequency distribution of extension lengths for each condition (three biological replicates for each condition). Addition of $Co^{2+}$ did not change the length distribution significantly.

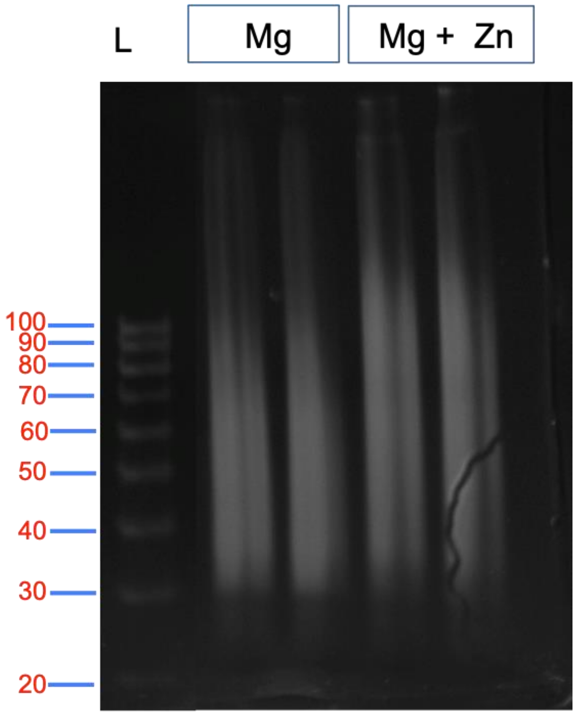


**Figure S4: Length distribution of extensions upon addition of $Zn^{2+}$ as seen on ssDNA gel**
Extension reactions were run as mentioned in Materials and Methods section. Two biological replicates per test condition were then loaded onto a ssDNA gel ($Mg^{2+}$ on left and $Mg^{2+}$+$Zn^{2+}$ on right). Addition of $Zn^{2+}$ increases the overall lengths of the extensions.

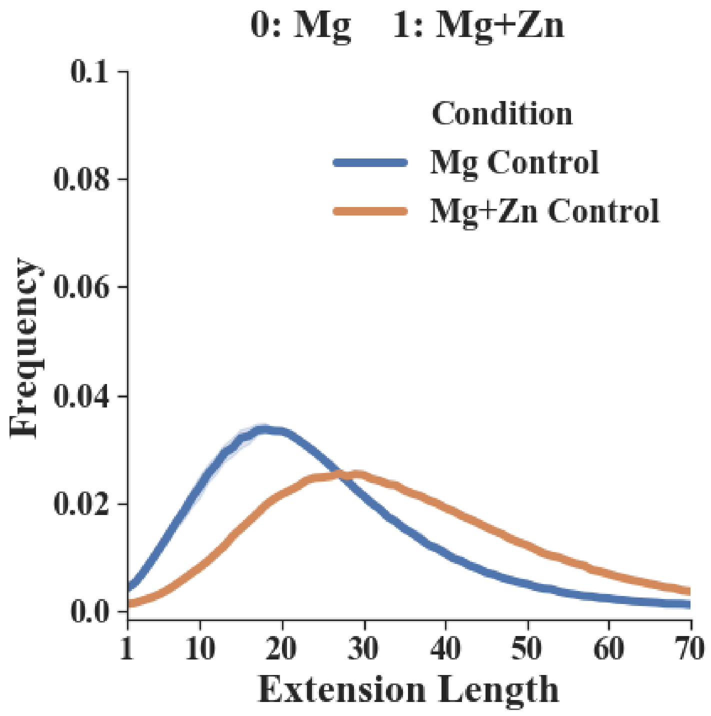


**Figure S5: Length distribution of extensions upon addition of $Zn^{2+}$ based on NGS data**
We calculated the mean frequency distribution of extension lengths for each condition (three biological replicates for each condition). Addition of $Zn^{2+}$ caused a shift in probability distribution toward longer lengths.

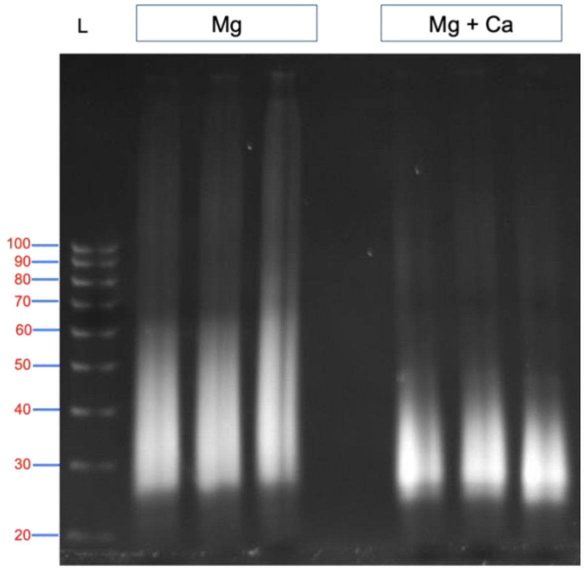


**Figure S6: Length distribution of extensions upon addition of $Ca^{2+}$ as seen on ssDNA gel**
Extension reactions were run as described in Materials and Methods section. Three biological replicates per test condition were then loaded on a ssDNA gel ($Mg^{2+}$ on left and $Mg^{2+}$+$Ca^{2+}$ on right). Addition of $Ca^{2+}$ decreases the overall lengths of the extensions.

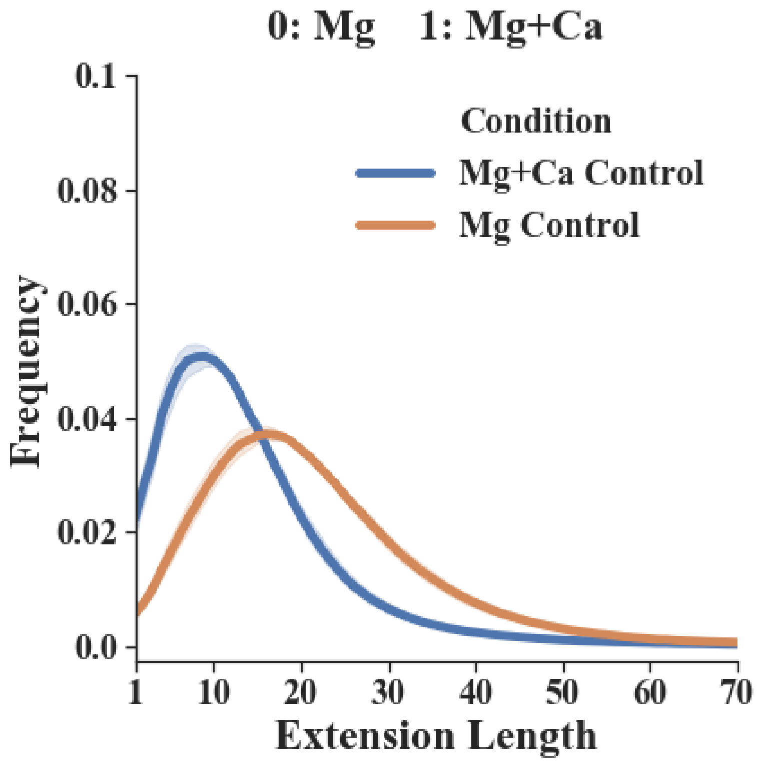


**Figure S7: Length distribution of extensions upon addition of $Ca^{2+}$ based on NGS data.**
We calculated the mean frequency distribution of extension lengths for each condition (three biological replicates for each condition). Addition of $Ca^{2+}$ caused a shift toward shorter lengths.

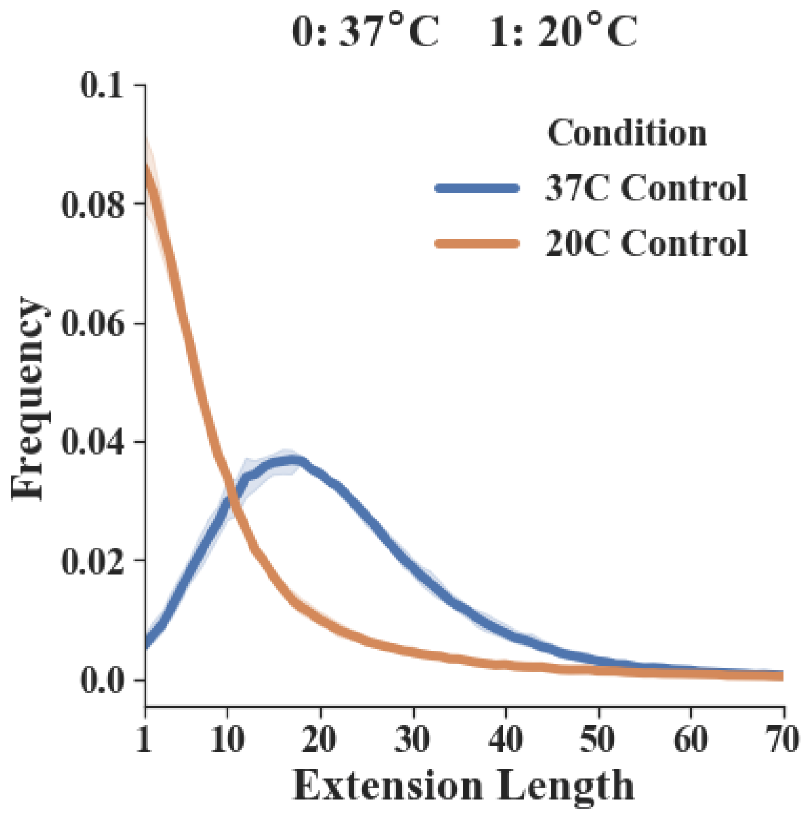


**Figure S8: Length distribution of extensions upon using temperature as a signal based on NGS data**
We calculated the mean frequency distribution of extension lengths for each condition (three biological replicates for each condition). Reducing the temperature of the extension reaction to 20 °C caused a shift toward shorter lengths.

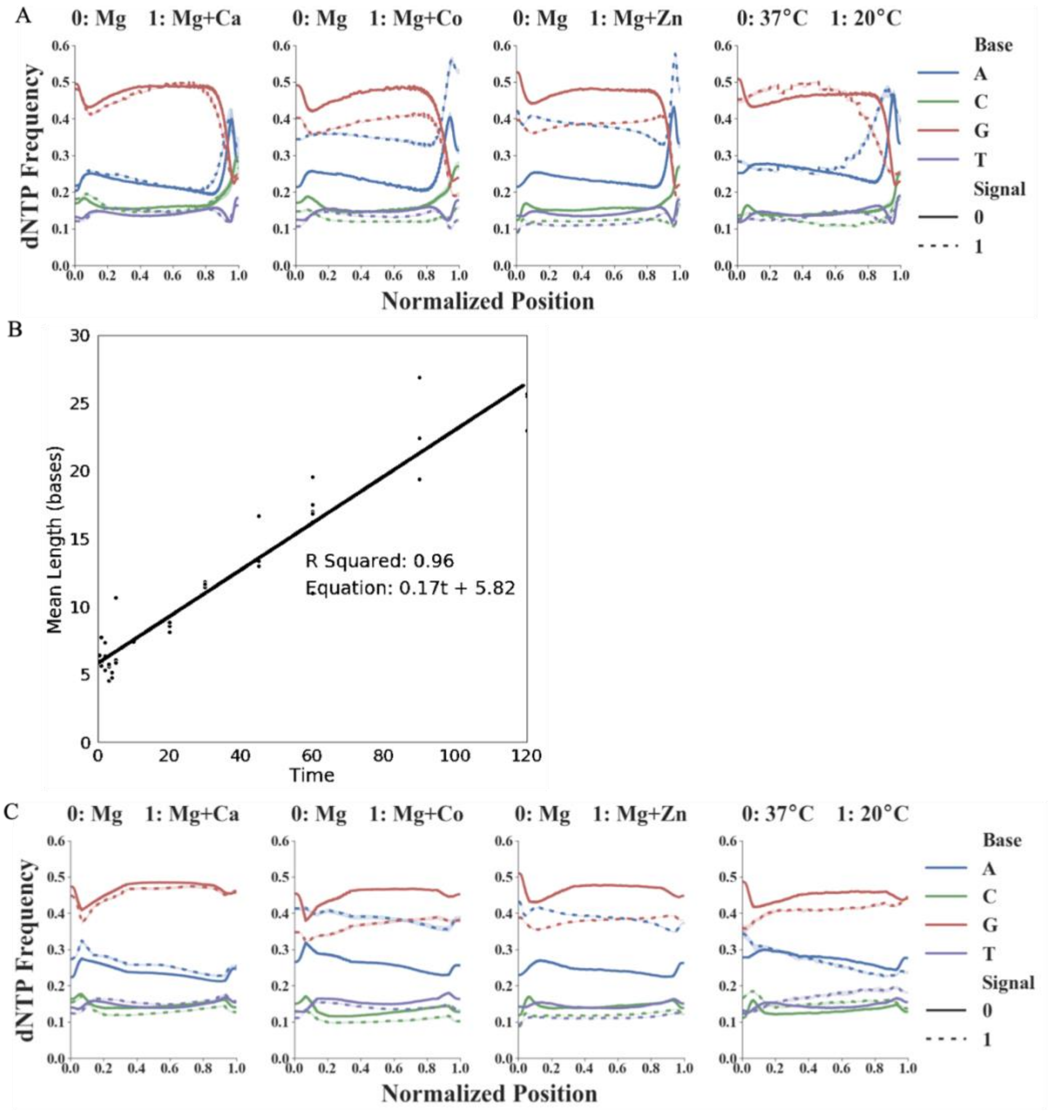


**Figure S9: Anomalous dNTP composition initially found at the end of reads and rate of reaction measured for extensions with only $Mg^{2+}$ present**

We observed a significant change in the individual dNTP frequency towards the ends of the ssDNA sequences synthesized. (A) Presents the significant change observed near the end of all reads with all the signals tested. Since we directly use 2 µL of extension reaction for ligation, the diluted TdT seems to be adding dNTPs to the ssDNA after the recording experiment, during the 16-hour ligation step. (B) To prove that these dNTPs were not added during the extension reaction (i.e. after the reaction), we sampled extension reactions (with $Mg^{2+}$ only) at several time points (Supplementary Text). We then calculated the mean extension length at each timepoint and applied a linear regression. The $R^2$ value of 0.96 for a straight line indicates that our assumption of constant rate (assuming input signal does not change) is valid. The slope of 0.17 reveals an average incorporation rate of 0.17 dNTPs/minute for this condition. Most importantly, the intercept of 5.82 indicates addition of 5.82 dNTPs (on average) either before or after the extension reaction. These are almost certainly being added after the extension reaction during the ligation step, which we conclude based on the anomalous behavior we see at the end of sequences in Panel A. (C) We created plots of the data from Panel A after trimming off last few dNTPs. See Materials and Methods for details on how these 5.8 bases were trimmed from the end of all sequences before further analysis.

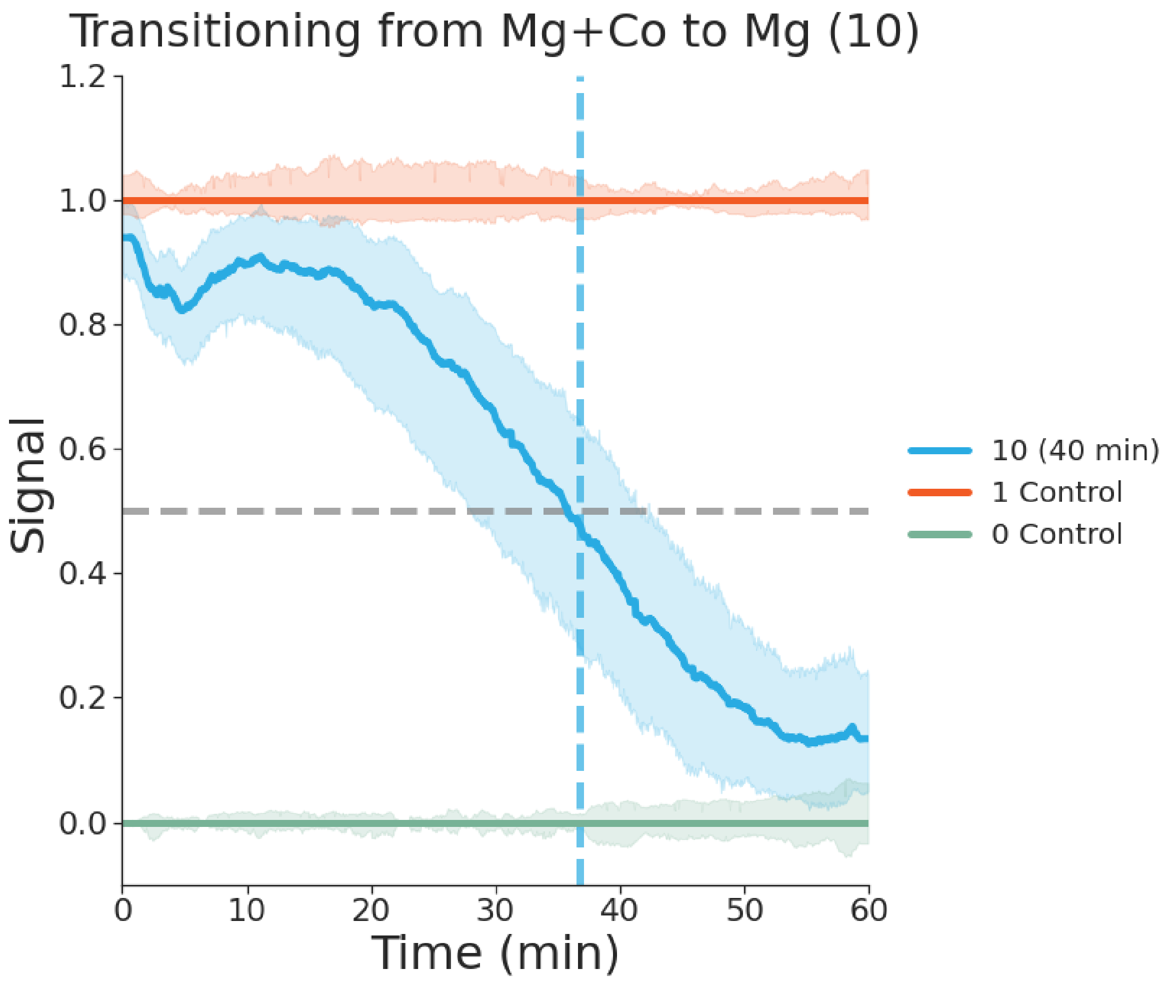


**Figure S10: Recording a single 1→0 step change in $Co^{2+}$ concentration onto ssDNA *in vitro.*** 0.25 mM $Co^{2+}$ was removed at 40 minutes to generate a 1→0 transition. This mean output signal across 6 biological replicates shows there is a difference in the preference of dNTP incorporated by TdT in the $Mg^{2+}$ (green) and $Mg^{2+}$+$Co^{2+}$ (orange) control conditions. The plot further shows the changes from 1→0 for $Co^{2+}$ removed at 40 minutes (blue). We were able to get a switch time of 36.7 minutes with a std. dev. of 9.9 minutes using methods used for 0→1 switch time predictions. We suspect the possible reason for higher variance in time prediction for this set-up was due to the ssDNA wash step at 40 minutes as discussed in Fig. S11.

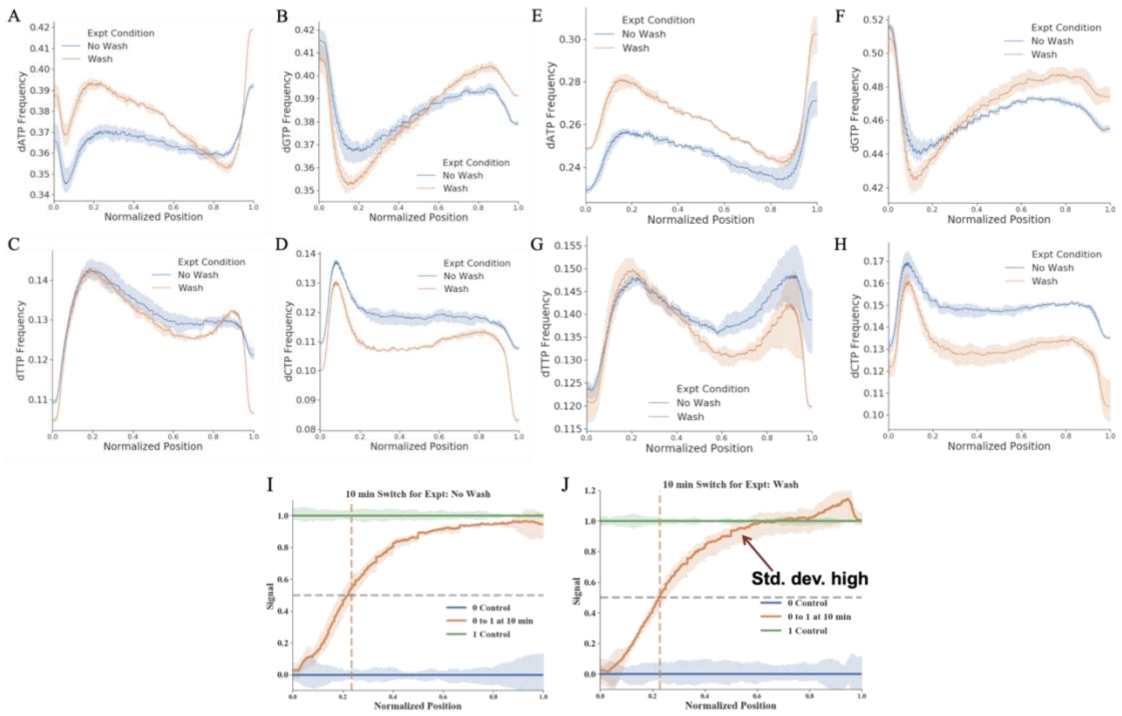


**Figure S11: dNTP bias & variability introduced by ssDNA wash columns**

We present here a comparison of the composition of sequences retained when the extension reactions were directly used for ligation (“No Wash”) vs. when the same extensions were put through a ssDNA wash kit (“Wash”). (A), (B), (C) and (D) show individual plots of each nucleotide frequency seen in extension reactions between No Wash vs Wash conditions for just $Mg^{2+}$ extensions. (E), (F), (G) and (H) show individual plots of each nucleotide frequency seen in extension reactions between No Wash vs Wash conditions for $Mg^{2+}$+ $Co^{2+}$ extensions. We observed a bias in overall dNTP content introduced by the columns used for ssDNA clean-up, when the reactions were washed after the recording experiment was stopped. ssDNA sequences with certain dNTP compositions were preferentially retained on the columns. (I) and (J) are plots for time prediction for No Wash and Wash condition respectively. We carried out an input signal of $Co^{2+}$ 0→1 at 10 minutes for a 1-hour extension. We obtained a time prediction of 12.8 minutes with 1.8 min std. dev. for No Wash condition. We obtained a time prediction of 12.4 min with a std. dev. of 1.2 min for the Wash condition. While the time predictions were very similar, there is a clear increase in variability (std. dev.) for the later part of the output signal recorded in (J) as compared to (I) (shown with a red arrow). Taken together, such biases and variability when introduced during the wash step for 0→1→0 experiment at 40 minutes for replacing +$Co^{2+}$ buffers with –$Co^{2+}$ buffers (See Materials and Methods: **Extension reactions for 0→1→0 set-up)** would cause more noise for the final 20 minutes of the recording.

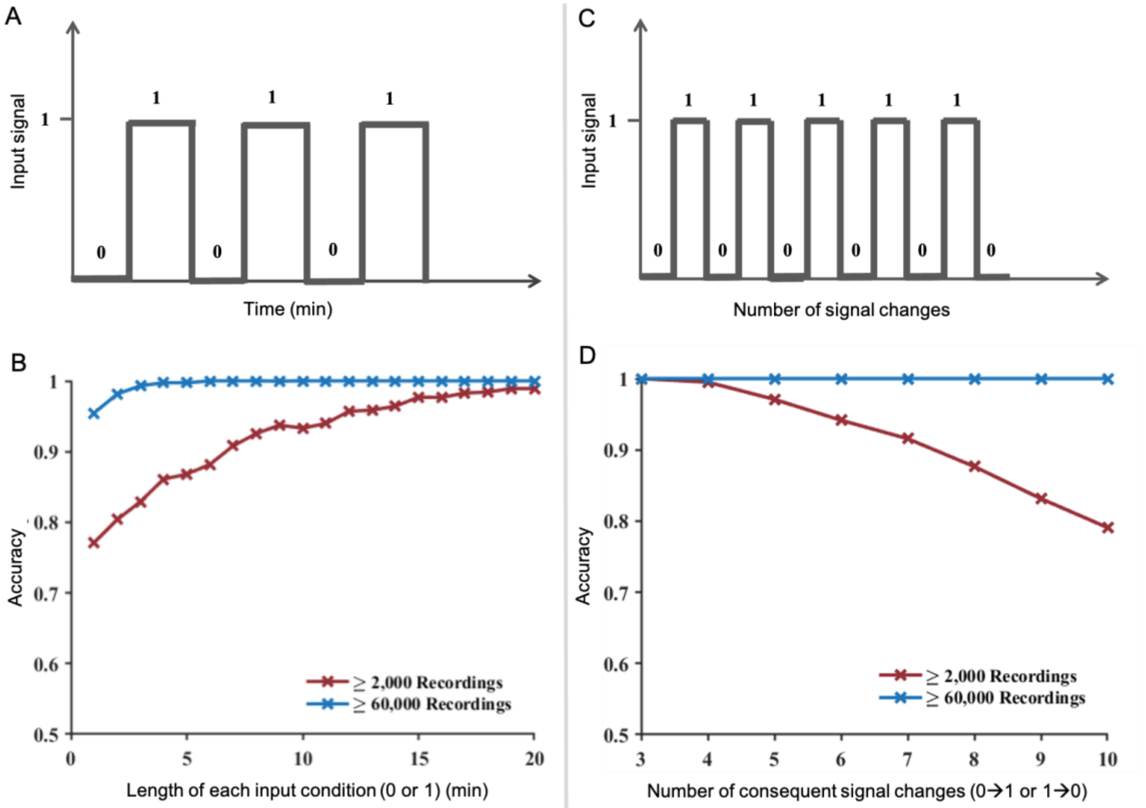


**Figure S12: *In silico* characterization of shortest resolvable input signal changes and highest number of consecutive signal changes that could be resolved using TURTLES**

*In silico* simulations based on experimental TdT parameters. For all simulations, we calculated the variability in two ways. The variability was either calculated across the first 100 nucleotides, in which there were at least 2000 recordings of all base numbers (red curve) or across the first 50 nucleotides, in which there were at least 60000 recordings of all base numbers (blue curve). (A) Representative signal for rapidly changing input from 0→1 or 1→0. (B) Output of *in silico* characterization of the signals represented by (A). We tried to find the shortest times of input changes from 0→1 or 1→0 that could be resolved using TURTLES. Simulations were carried out where the number of input signal changes were kept constant at 6 (e.g., 0→1→0→1→0→1→0) while the length of time for each input condition (0 or 1) was varied. Even with just 2000 strands of ssDNA recorded (each 100bp in length), input signal changing every 1 minute was resolved with > 75% accuracy, and input signal changing every 10 minutes was resolved with > 90% accuracy. (C) Representative signal for multiple changes in input signal from 0→1 or 1→0. (D) Output of *in silico* characterization of the signals represented by (C). Simulations were carried out with the duration of each input condition (0 or 1) kept constant at 10 minutes while varying the number of input signal changes. The accuracy of resolving this output signal was calculated across simulations. Even with just 2000 strands of ssDNA recorded (each 100bp in length), 3 input signal changes could be resolved with almost 100% accuracy and 10 input signal changes of 10 minutes each could be resolved at about 80% accuracy.

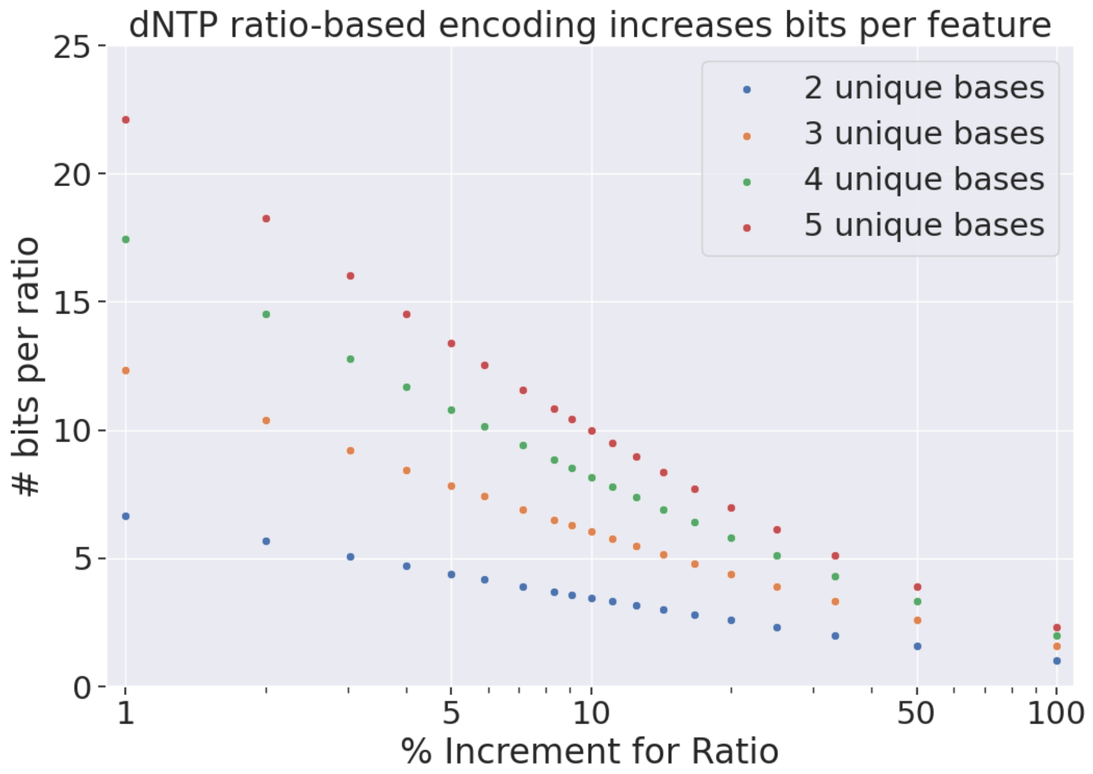


**Figure S13:** Storage potential with a TdT-based system encoding data using nucleotide composition rather than sequence identity. For example, with 4 unique bases (A, C, G, and T) and a set of ratios built in increments of 5% (e.g. 35% A, 20% C, 15% G, 30% T), the maximum amount of data stored per ratio is 11 bits. Even greater densities are possible with smaller increments or use of unnatural bases.

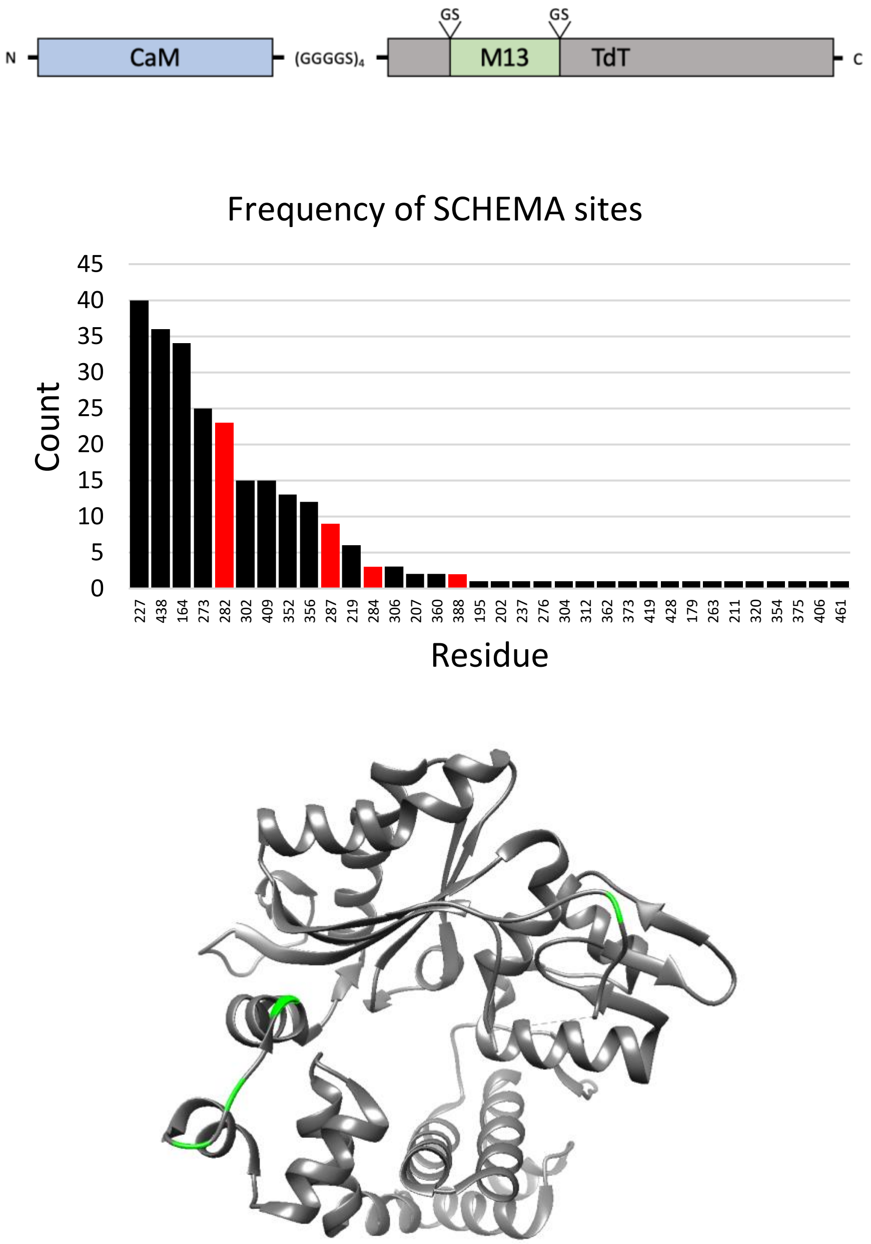


**Figure S14: CaM-TdT(M13) design and SCHEMA crossover site frequency and locations.** (A) schematic representation of CaM-TdT(M13) variants. (B) the number of times each crossover site appeared in the SCHEMA/RASPP analysis. Sites selected for fusions are colored in red. (C) ribbon diagram of the catalytic core of the short isoform of murine TdT (PDB ID: 4i27) with fusion site residues colored in green. Fusion sites were selected to minimize steric interference and target polymerase regions required for DNA binding (282,284,287) or catalysis (388).

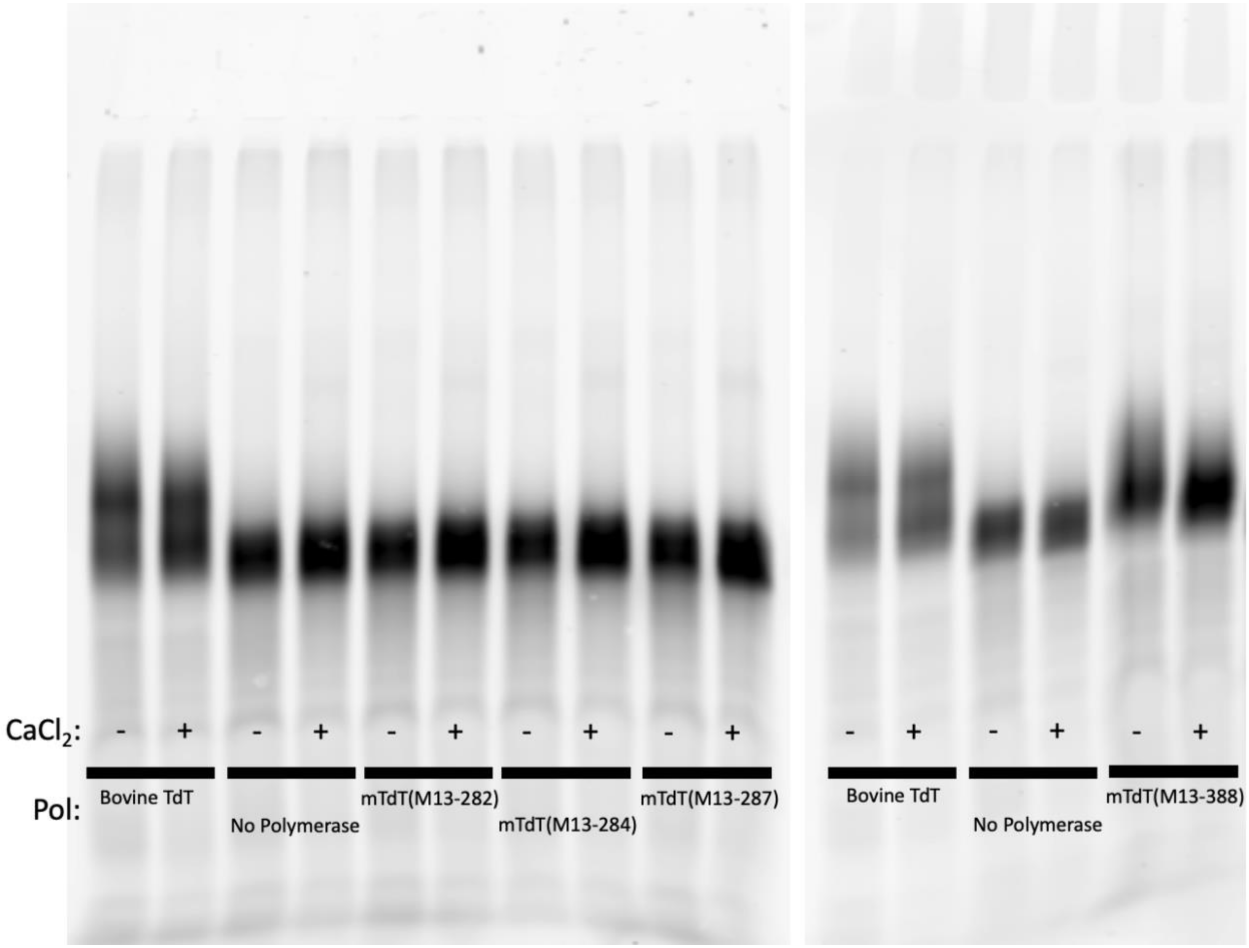


**Figure S15: Primer extension activity of M13 TdT fusion proteins.** Fusion proteins were screened by assaying the activity of the M13 fusion variants without N-terminal CaM to ensure that mTdT remained active with the M13 fusion. Fusions were expressed in PURExpress and their activity was screened by primer extension. Primer extension reactions were performed without added $CaCl_2$ (indicated by "-") and with 1mM $CaCl_2$ (indicated by "+"). The reaction products were visualized by urea-PAGE (see methods). Fusions are labelled as mTdT(M13-###), where the number indicates the residue immediately preceding the GS-M13-GS fusion.

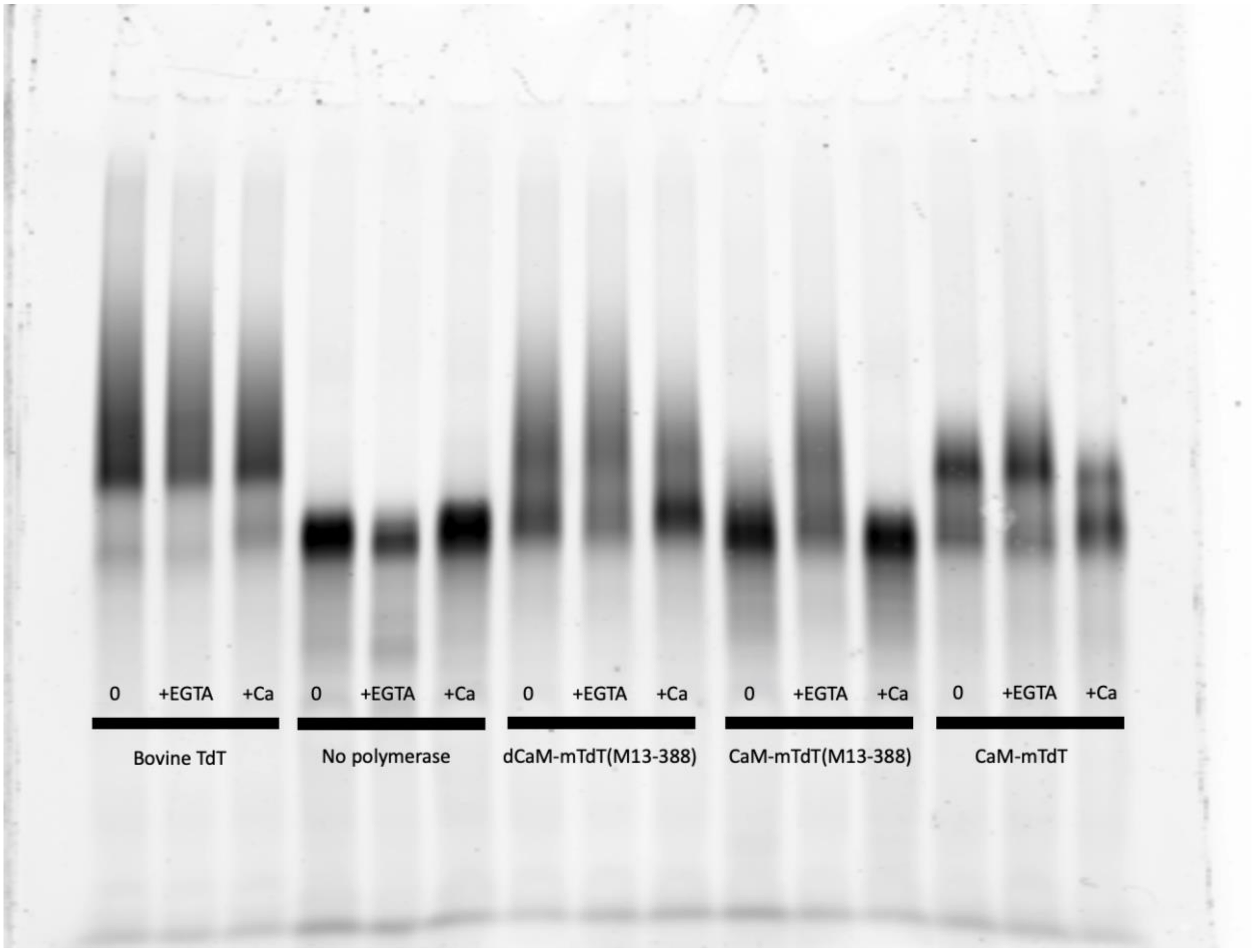


**Figure S16: Primer extension activity of CaM-mTdT(M13-388), dCaM-mTdT(M13-388), and mTdT(M13-388).** The activity of each polymerase was tested in three conditions where "0" indicates reactions with no added calcium or EGTA, "+EGTA" indicates reactions supplemented with 1 mM EGTA, and "+Ca" indicates reactions supplemented with 1 mM CaCl2. dCaM-mTdT(M13-388) contains CaM mutations D20A, D56A, D93A, and D129A that ablate the calcium-binding capability of CaM(49). CaM-mTdT consists only of mTdT with N-terminal CaM fusion.

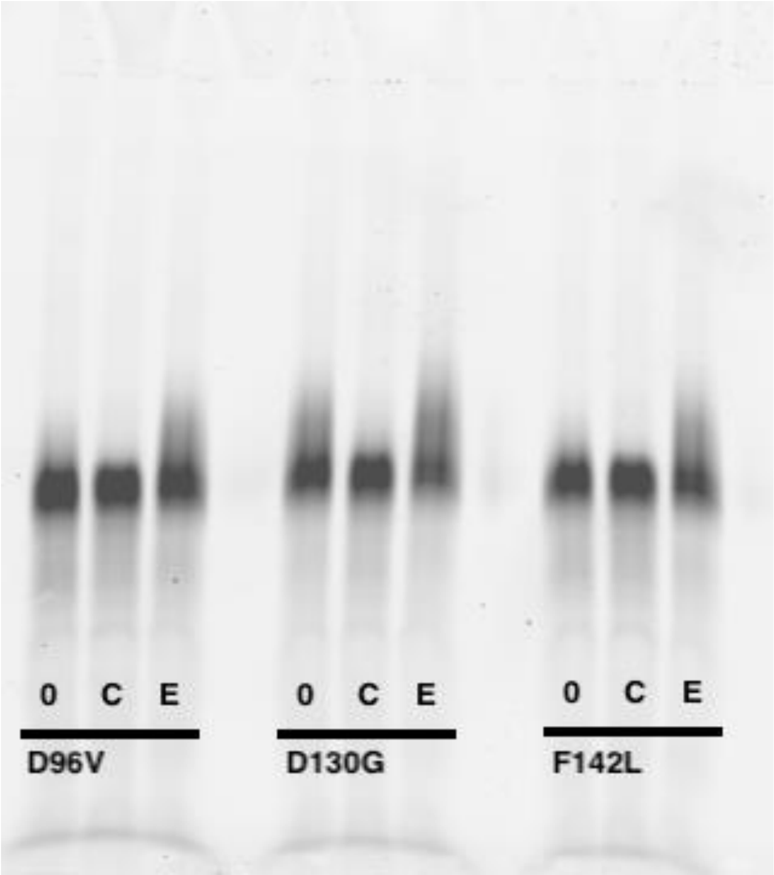


**Figure S17: Primer extension activity of CaM affinity variants.** the activity of calcium affinity variants was tested by primer extension and visualized by urea PAGE in non-supplemented (“0”), 1 mM $CaCl_2$ added (“C”), and 1mM EGTA added conditions (“E”). Crotti *et al.* report effective calcium affinities of 38 μM, 150 μM, and 15μM respectively as compared to 2.8μM for wildtype CaM.

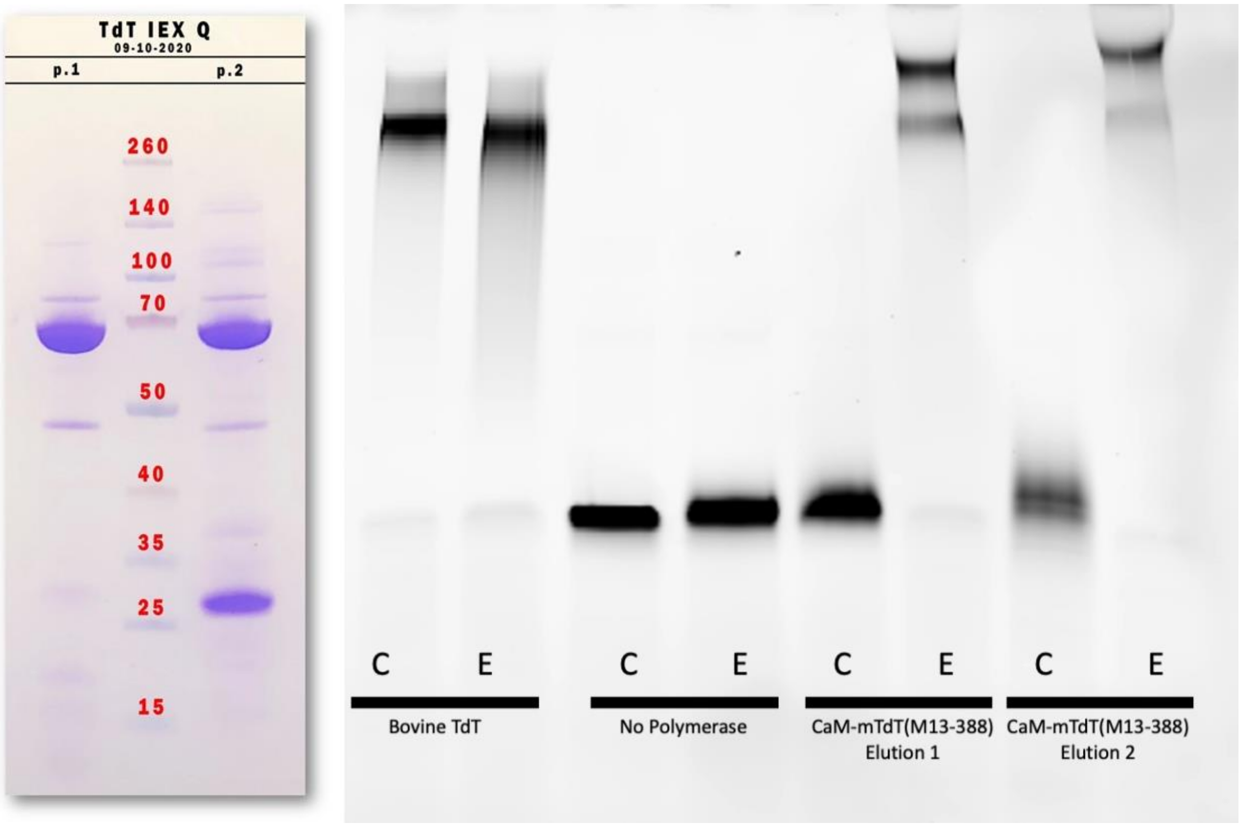


**Figure S18: Primer extension activity of purified CaM-mTdT(M13-388). A)** Anion exchange elutions 1 (p.1) and 2 (p.2), visualized by PAGE. CaM-mTdT(M13-388) appears at approximately 70kDa. (B) Following buffer exchange, the activity of both fractions was assayed by primer extension and compared to the wildtype polymerase and MBP-CaM-mTdT(M13-388) (from PURExpress). 1 mM $CaCl_2$ added (“C”), and 1mM EGTA added conditions (“E”).

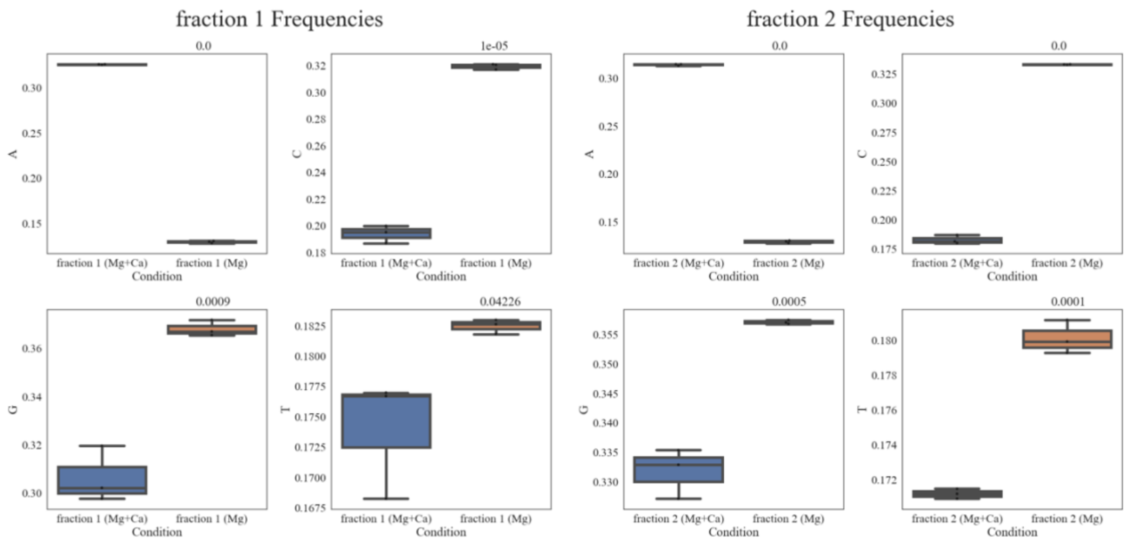


**Figure S19.  Comparison of purified CaM-mTdT(M13-388) fractions.** base incorporation frequencies of CaM-mTdT(m13-388) elution fractions 1 and 2 in primer extension reactions.

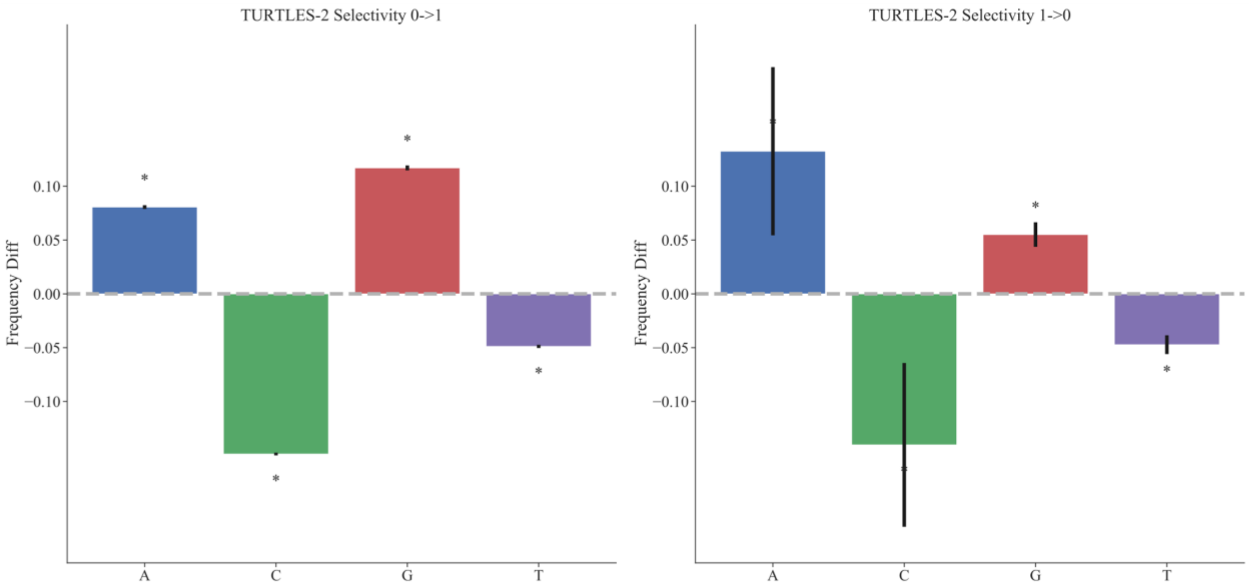


**Figure S20: Incorporation preference of two-polymerase recording system.** The difference in base selectivity of TURTLES-2 in the presence of calcium as compared to calcium-free conditions.

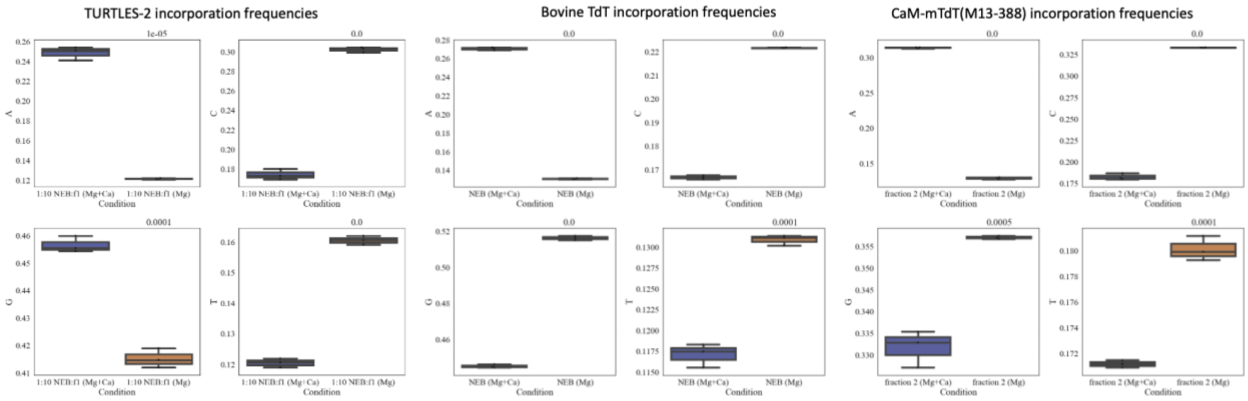


**Figure S21: TURTLE-2 Selectivity comparison.** Incorporation frequencies of each base for the two-polymerase TURTLES-2 system, bovine TdT, and purified CaM-mTdT(M13-388).

### Supplementary Tables

| Name | Sequence | Notes |
|---|---|---|
| CS1 | ACACTGACGACATGGTTCTACA | |
| CS1_5N | ACACTGACGACATGGTTCTACANNNNN | Degenerate "N " bases are distributed 25% A, 15%C , 45%G, 15% T |
| FAM_NB | (6'-FAM)A*C*A*C*TGACGACATGGTTCTACAACCGGTATGTACGGCGGTCGGTTATCGTA | "*" follows phosphorothioated bases |
| AMD006 | AGGCTAGTCGTCTGTATAGG | |
| P1 | CTCCAGAGGCCACAGGGTCTGCTGACCAATTAACCGAAGAAC | |
| P2 | TGTTCCTCGTCCTCGCTGCCCTTGGCTGTCATCATCTGC | |
| P3 | AGGCCACAGAGGACGGGTCTGCTGACCAATTAACCGAAGAAC | |
| P4 | AACTGTTGTTCCTCGCTGCCCTTGGCTGTCATCATCTGC | |
| P5 | CCTTCGAGAAGTTCGGGTCTGCTGACCAATTAACCGAAGAAC | |
| P6 | CGGCTTGGCTGTTTGCTGCCCTTGGCTGTCATCATCTGC | |
| P7 | AAGGTAAAGGCTGGGGGTCTGCTGACCAATTAACCGAAGAAC | |
| P8 | ACACGGATAGCTTTGCTGCCCTTGGCTGTCATCATCTGC | |
| P9 | AATTCGACGACGACGACAAAGCTGACCAATTAACCGAAGAACAAATC | |
| P10 | TGCGAAATCTTTTTTACCGCGGAACCTCCCCCTCCAGAAC | |
| P11 | TGATGACAGCCAAGGGCAGCGAGGACGAGGAACAACAGTTG | |
| P12 | GTTAATTGGTCAGCAGACCCTGTGGCCTCTGGAGAAGTGATC | |
| P13 | TGATGACAGCCAAGGGCAGCGAGGAACAACAGTTGTTGCACAAG | |
| P14 | GTTAATTGGTCAGCAGACCCGTCCTCTGTGGCCTCTGGAG | |
| P15 | TGATGACAGCCAAGGGCAGCAAACAGCCAAGCCGCAAAG | |
| P16 | GTTAATTGGTCAGCAGACCCGAACTTCTCGAAGGTACTTTCTAAAATATCGC | |
| P17 | TGATGACAGCCAAGGGCAGCAAAGCTATCCGTGTTGACCTTGTG | |
| P18 | GTTAATTGGTCAGCAGACCCCCAGCCTTTACCTTCCTGCTG | |
| P19 | GTTCTGGAGGGGGAGGTTCCGCGGTAAAAAAGATTTCGCAGTACG | |
| P20 | TCAGCTTTGTCGTCGTCGTCGAATTCAGACCCGTCGACGATG | |
| P21 | AGATCCAAAGTGACAAAAGCGGGTCTGCACGCCGTAAATG | |
| P22 | TGCATTTGCGTGAAACGCAGGCTGCCGGAACTTAAACGCC | |
| P23 | AAAGTGACAAAAGCCTGCGTGGGTCTGCACGCCGTAAATG | |
| P24 | GCTTTTTGCATTTGCGTGAAGCTGCCGGAACTTAAACGCC | |
| P25 | AAAGCCTGCGTTTCACGCAAGGGTCTGCACGCCGTAAATG | |
| P26 | AAAAACCCAGCTTTTTGCATGCTGCCGGAACTTAAACGCC | |
| P27 | AAAGTACCTTCGAGAAGTTCGGGTCTGCACGCCGTAAATG | |
| P28 | ACTTTGCGGCTTGGCTGTTTGCTGCCGGAACTTAAACGCC | |
| P29 | AGATCCAAAGTGACAAAAGCGGGTCTGCACGCCGTAAATG | |
| P30 | TGCATTTGCGTGAAACGCAGGCTGCCGGAACTTAAACGCC | |
| P31 | AAAGTGACAAAAGCCTGCGTGGGTCTGCACGCCGTAAATG | |
| P32 | GCTTTTTGCATTTGCGTGAAGCTGCCGGAACTTAAACGCC | |
| P33 | AAAGCCTGCGTTTCACGCAAGGGTCTGCACGCCGTAAATG | |
| P34 | AAAAACCCAGCTTTTTGCATGCTGCCGGAACTTAAACGCC | |
| P35 | AAAGTACCTTCGAGAAGTTCGGGTCTGCACGCCGTAAATG | |
| P36 | ACTTTGCGGCTTGGCTGTTTGCTGCCGGAACTTAAACGCC | |

| | |
|---|---|
| S1 | GCATTCTGGTATGCCGTCCG |
| S2 | GGTGGACGAAATGATCCGCG |
| S3 | GTTCGTGTGTCGCGTAGCGG |
| S4 | GGCTGAAGCGGTAAGTATGTTGG |
| S5 | CGAGGAGGAAATCCGCGAAGC |
| S6 | GATCACTTCTCCAGAGGCCACAG |
| S7 | GAGCCTTCCATTCCCTATTACCAG |
| S8 | GTCCGGCGTAGAGGATCGAG |
| S9 | CACTTGACAAGGAACTGAAGGCC |

**Table S1: DNA substrate sequences and primer sequences**

| Name | Amino Acid Sequence |
| --- | --- |
| CaM-mTdT(M13-388) | MGHHHHHHHHHHSSGHIDDDDKHADQLTEEQIAEFKEAFSLFDKDGDGTITTKELGTVMRSLGQNPTEAELQDMINEVDADGNGTIDFPEFLTMMARKMKDTDSEEEIREAFRVFDKDGNGYISAAELRHVMTNLGEKLTDEEVDEMIREADID<br>GDGQVNYEEFVQMMTAKGGGGSGGGGSGGGGSGGGGSAVKKISQYACQRRTTLNNYNQLFTDALDILAENDELRENEGSCLAFMRASSVLKSLPFPITSMKDTEGIPCLGDKVKSIIEGIIEDGESSEAKAVLNDERYKSFKLFTSVFGVGLKT<br>AEKWFRMGFRTLSKIQSDKSLRFTQMQKAGFLYYEDLVSCVNRPEAEAVSMLVKEAVVTFLPDALVTMTGGFRRGKMTGHDVDFLITSPEATEDEEQQLLHKVTDFWKQQGLLLYCDILESTFEKFGSARRKWQKTGHAVRAIGRLSSGSKQPS<br>RKVDALDHFQKCFLILKLDHGRVHSEKSGQQEGKGWKAIRVDLVMCPYDRRAFALLGWTGSRQFERDLRRYATHERKMMLDNHALYDRTKRVFLEAESEEEIFAHLGLDYIEPWERNA |
| MBP-CaM-mTdT(M13-282) | MGHHHHHHHHHHSSGHIDDDDKHMMKIEEGKLVIWINGDKGYNGLAEVGKKFEKDTGIKVTVEHPDKLEEKFPQVAATGDGPDIIFWAHDRFGGYAQSGLLAEITPDKAFQDKLYPFTWDAVRYNGKLIAYPIAVEALSLIYNKDLLPNPPKTW<br>EEIPALDKELKAKGKSALMFNLQEPYFTWPLIAADGGYAFKYENGKYDIKDVGVDNAGAKAGLTFLVDLIKNKHMNADTDYSIAEAAFNKGETAMTINGPWAWSNIDTSKVNYGVTVLPTFKGQPSKPFVGVLSAGINAASPNKELAKEFLENY<br>LLTDEGLEAVNKDKPLGAVALKSYEEELVKDPRIAATMENAQKGEIMPNIPQMSAFWYAVRTAVINAASGRQTVDEALKDAQTNSSSNNNNNNNNNNLGIEGRISHMSMGGRDIVDGSEFDDDDKADQLTEEQIAEFKEAFSLFDKDGDGTITT<br>KELGTVMRSLGQNPTEAELQDMINEVDADGNGTIDFPEFLTMMARKMKDTDSEEEIREAFRVFDKDGNGYISAAELRHVMTNLGEKLTDEEVDEMIREADIDGDGQVNYEEFVQMMTAKGGGGSGGGGSGGGGSGGGGSAVKKISQYACQRRTT<br>LNNYNQLFTDALDILAENDELRENEGSCLAFMRASSVLKSLPFPITSMKDTEGIPCLGDKVKSIIEGIIEDGESSEAKAVLNDERYKSFKLFTSVFGVGLKTAEKWFRMGFRTLSKIQSDKSGSARRKWQKTGHAVRAIGRLSSGSLRFTQMQK<br>AGFLYYEDLVSCVNRPEAEAVSMLVKEAVVTFLPDALVTMTGGFRRGKMTGHDVDFLITSPEATEDEEQQLLHKVTDFWKQQGLLLYCDILESTFEKFKQPSRKVDALDHFQKCFLILKLDHGRVHSEKSGQQEGKGWKAIRVDLVMCPYDRRA<br>FALLGWTGSRQFERDLRRYATHERKMMLDNHALYDRTKRVFLEAESEEEIFAHLGLDYIEPWERNA |
| MBP-CaM-mTdT(M13-284) | MGHHHHHHHHHHSSGHIDDDDKHMMKIEEGKLVIWINGDKGYNGLAEVGKKFEKDTGIKVTVEHPDKLEEKFPQVAATGDGPDIIFWAHDRFGGYAQSGLLAEITPDKAFQDKLYPFTWDAVRYNGKLIAYPIAVEALSLIYNKDLLPNPPKTW<br>EEIPALDKELKAKGKSALMFNLQEPYFTWPLIAADGGYAFKYENGKYDIKDVGVDNAGAKAGLTFLVDLIKNKHMNADTDYSIAEAAFNKGETAMTINGPWAWSNIDTSKVNYGVTVLPTFKGQPSKPFVGVLSAGINAASPNKELAKEFLENY<br>LLTDEGLEAVNKDKPLGAVALKSYEEELVKDPRIAATMENAQKGEIMPNIPQMSAFWYAVRTAVINAASGRQTVDEALKDAQTNSSSNNNNNNNNNNLGIEGRISHMSMGGRDIVDGSEFDDDDKADQLTEEQIAEFKEAFSLFDKDGDGTITT<br>KELGTVMRSLGQNPTEAELQDMINEVDADGNGTIDFPEFLTMMARKMKDTDSEEEIREAFRVFDKDGNGYISAAELRHVMTNLGEKLTDEEVDEMIREADIDGDGQVNYEEFVQMMTAKGGGGSGGGGSGGGGSGGGGSAVKKISQYACQRRTT<br>LNNYNQLFTDALDILAENDELRENEGSCLAFMRASSVLKSLPFPITSMKDTEGIPCLGDKVKSIIEGIIEDGESSEAKAVLNDERYKSFKLFTSVFGVGLKTAEKWFRMGFRTLSKIQSDKSLRGSARRKWQKTGHAVRAIGRLSSGSFTQMQK<br>AGFLYYEDLVSCVNRPEAEAVSMLVKEAVVTFLPDALVTMTGGFRRGKMTGHDVDFLITSPEATEDEEQQLLHKVTDFWKQQGLLLYCDILESTFEKFKQPSRKVDALDHFQKCFLILKLDHGRVHSEKSGQQEGKGWKAIRVDLVMCPYDRRA<br>FALLGWTGSRQFERDLRRYATHERKMMLDNHALYDRTKRVFLEAESEEEIFAHLGLDYIEPWERNA |
| MBP-CaM-mTdT(M13-287) | MGHHHHHHHHHHSSGHIDDDDKHMMKIEEGKLVIWINGDKGYNGLAEVGKKFEKDTGIKVTVEHPDKLEEKFPQVAATGDGPDIIFWAHDRFGGYAQSGLLAEITPDKAFQDKLYPFTWDAVRYNGKLIAYPIAVEALSLIYNKDLLPNPPKTW<br>EEIPALDKELKAKGKSALMFNLQEPYFTWPLIAADGGYAFKYENGKYDIKDVGVDNAGAKAGLTFLVDLIKNKHMNADTDYSIAEAAFNKGETAMTINGPWAWSNIDTSKVNYGVTVLPTFKGQPSKPFVGVLSAGINAASPNKELAKEFLENY<br>LLTDEGLEAVNKDKPLGAVALKSYEEELVKDPRIAATMENAQKGEIMPNIPQMSAFWYAVRTAVINAASGRQTVDEALKDAQTNSSSNNNNNNNNNNLGIEGRISHMSMGGRDIVDGSEFDDDDKADQLTEEQIAEFKEAFSLFDKDGDGTITT<br>KELGTVMRSLGQNPTEAELQDMINEVDADGNGTIDFPEFLTMMARKMKDTDSEEEIREAFRVFDKDGNGYISAAELRHVMTNLGEKLTDEEVDEMIREADIDGDGQVNYEEFVQMMTAKGGGGSGGGGSGGGGSGGGGSAVKKISQYACQRRTT<br>LNNYNQLFTDALDILAENDELRENEGSCLAFMRASSVLKSLPFPITSMKDTEGIPCLGDKVKSIIEGIIEDGESSEAKAVLNDERYKSFKLFTSVFGVGLKTAEKWFRMGFRTLSKIQSDKSLRFTQGSARRKWQKTGHAVRAIGRLSSGSMQK<br>AGFLYYEDLVSCVNRPEAEAVSMLVKEAVVTFLPDALVTMTGGFRRGKMTGHDVDFLITSPEATEDEEQQLLHKVTDFWKQQGLLLYCDILESTFEKFKQPSRKVDALDHFQKCFLILKLDHGRVHSEKSGQQEGKGWKAIRVDLVMCPYDRRA<br>FALLGWTGSRQFERDLRRYATHERKMMLDNHALYDRTKRVFLEAESEEEIFAHLGLDYIEPWERNA |
| MBP-CaM-mTdT(M13-388) | MGHHHHHHHHHHSSGHIDDDDKHMMKIEEGKLVIWINGDKGYNGLAEVGKKFEKDTGIKVTVEHPDKLEEKFPQVAATGDGPDIIFWAHDRFGGYAQSGLLAEITPDKAFQDKLYPFTWDAVRYNGKLIAYPIAVEALSLIYNKDLLPNPPKTW<br>EEIPALDKELKAKGKSALMFNLQEPYFTWPLIAADGGYAFKYENGKYDIKDVGVDNAGAKAGLTFLVDLIKNKHMNADTDYSIAEAAFNKGETAMTINGPWAWSNIDTSKVNYGVTVLPTFKGQPSKPFVGVLSAGINAASPNKELAKEFLENY<br>LLTDEGLEAVNKDKPLGAVALKSYEEELVKDPRIAATMENAQKGEIMPNIPQMSAFWYAVRTAVINAASGRQTVDEALKDAQTNSSSNNNNNNNNNNLGIEGRISHMSMGGRDIVDGSEFDDDDKADQLTEEQIAEFKEAFSLFDKDGDGTITT<br>KELGTVMRSLGQNPTEAELQDMINEVDADGNGTIDFPEFLTMMARKMKDTDSEEEIREAFRVFDKDGNGYISAAELRHVMTNLGEKLTDEEVDEMIREADIDGDGQVNYEEFVQMMTAKGGGGSGGGGSGGGGSGGGGSAVKKISQYACQRRTT<br>LNNYNQLFTDALDILAENDELRENEGSCLAFMRASSVLKSLPFPITSMKDTEGIPCLGDKVKSIIEGIIEDGESSEAKAVLNDERYKSFKLFTSVFGVGLKTAEKWFRMGFRTLSKIQSDKSLRFTQMQKAGFLYYEDLVSCVNRPEAEAVSML<br>VKEAVVTFLPDALVTMTGGFRRGKMTGHDVDFLITSPEATEDEEQQLLHKVTDFWKQQGLLLYCDILESTFEKFGSARRKWQKTGHAVRAIGRLSSGSKQPSRKVDALDHFQKCFLILKLDHGRVHSEKSGQQEGKGWKAIRVDLVMCPYDRRA<br>FALLGWTGSRQFERDLRRYATHERKMMLDNHALYDRTKRVFLEAESEEEIFAHLGLDYIEPWERNA |

**Table S2: Amino acid sequences of TdT-CaM fusions**

Expression vector sequences available upon request
